# A self-supervised deep learning pipeline for segmentation in two-photon fluorescence microscopy

**DOI:** 10.1101/2025.01.20.633744

**Authors:** Emmanuel Edward Ntiri, Tony Xu, Matthew Rozak, Ahmadreza Attarpour, Adrienne Dorr, Bojana Stefanovic, Maged Goubran

## Abstract

Two-photon fluorescence microscopy (TPFM) allows in situ investigation of the structure and function of the brain at a cellular level, but the conventional image analyses of TPFM data are labour-intensive. Automated deep learning (DL)-based image processing pipelines used to analyze TPFM data require large labeled training datasets. Here, we developed a self supervised learning (SSL) pipeline to test whether unlabeled data can be used to boost the accuracy and generalizability of DL models for image segmentation in TPFM. We specifically developed four pretext tasks, including shuffling, rotation, axis rotation, and reconstruction, to train models without supervision using the UNet architecture. We validated our pipeline on two tasks (neuronal soma and vasculature segmentation), using large 3D microscopy datasets. We introduced a novel density-based metric, which provided more sensitive evaluation to downstream analysis tasks. We further determined the amount of labeled data required to reach performance on par with fully supervised learning (FSL) models. SSL-based models that were fine-tuned with only 50% of data were on par or superior (e.g., Dice increase of 3% for neuron segmentation and Dice score of 0.88 ± 0.09 for vessel segmentation) to FSL models. We demonstrated that segmentation maps generated by SSL models pretrained on the reconstruction and rotation tasks can be better translated to downstream tasks than can other SSL tasks. Finally, we benchmarked all models on a publicly available out-of-distribution dataset, demonstrating that SSL models outperform FSL when trained with clean data, and are more robust than FSL models when trained with noisy data.

## Introduction

Two-photon fluorescence microscopy (TPFM) enables *in situ* cellular-scale exploration of the microscopic structure and function of the brain. Segmentation and classification of objects is an important analysis step in TPFM analyses, as the volume and morphological properties of anatomical structures may provide insights into disease state and progression (Bennett et al., 2018; Rozak et al., 2024; Tsai et al., 2009). The advent of deep learning (DL) has led to a push for the creation of tools to address segmentation as a bottleneck in TPFM analysis pipelines.

Convolutional neural networks (CNNs) have been applied to segmentation tasks in multi-photon microscopy and light-sheet fluorescent microscopy (Cai et al., 2020; Haft-Javaherian et al., 2019; Klinghoffer et al., 2020a; Todorov et al., 2020); however, they are usually tailored to a specific structure, and often generalize poorly to out-of-distribution data (Ciga et al., 2022; Krug and Rohr, 2022; Srinidhi et al., 2022). State-of-the-art (SOTA) models such as the commonly used UNet architecture (Ronneberger et al., 2015) require large datasets to achieve robust performances. Generating ground truth annotations in 3D microscopy is exceptionally time-consuming, suffers from interrater variability, and requires data-specific expertise.

Self supervised learning (SSL) may serve as an avenue to leverage datasets without annotations to improve the efficiency and generalizability of image segmentation in microscopy. SSL uses unlabeled data to improve performance on downstream tasks (Gui et al., 2023) as models may learn semantic representations of the data without prompting from labels (Srinidhi et al., 2022). Training models with SSL may be of great benefit for TPFM studies, where there is an abundance of large image volumes and datasets. There are two main tasks in SSL schemes: a pretext task and a downstream task. In the pretext (or auxiliary) task, pseudo labels can be generated from unlabeled data, to learn representations of the data (Chaitanya et al., 2020).

The weights from the initial task are then transferred to the downstream task, and are finetuned with labeled data. SSL approaches have been previously applied in microscopy. SSL-based methods have been applied on single to few-shot downstream tasks, where pretrained models are immediately tested on downstream tasks with little or no finetuning, in microscopy datasets (Dawoud et al., 2022; Midtvedt et al., 2022; Ouyang et al., 2022). As a pretext task, Krug and Rohr (Krug and Rohr, 2022), trained models to classify the magnification scale of images in 2D epi-fluorescence microscopy videos. Pretrained models were then finetuned to segment cells in fluorescent microscopy datasets. Lu et al. (Lu et al., 2019) leveraged the multiple channels in fluorescent microscopy datasets in an “inpainting” task, where models were tasked with predicting the fluorescent pattern in a second image given the cell image of the first. While SSL applications have been extended to 3D TPFM data, they have been applied only to the downstream tasks of image denoising and super-resolution (He et al., 2023; Wang et al., 2021).

To explore the impact of SSL when applied to limited amounts of labelled TPFM data, we developed SELF-TPFM: a Pytorch-based pipeline to train and test SSL models for the segmentation of microscopy data. Our pipeline uses several types of pretext tasks: shuffling, rotation, rotation along an axis, and reconstruction; and includes methods for pretraining, finetuning, and testing SSL models with existing datasets. To our knowledge, our pipeline is the first to investigate the effectiveness of different pretext tasks against fully supervised learning (FSL) in TPFM. To enhance model evaluation, we introduce a novel density-based metric that computes the density of foreground voxels within image patches, which provides more sensitive evaluation to downstream analysis tasks than standard segmentation overlap metrics. By comparing CNN-based models trained using SSL to FSL models trained with varying degrees of labeled neuron and vessel TPFM datasets, we demonstrate that pretrained models with SSL segment neurons and vessels as accurately as FSL models trained with the same amount of data. Furthermore, we found that models trained on the reconstruction-based pretext task outperform FSL models by 3% when finetuned with half the amount of neuronal data. We perform additional experiments to assess the OOD generalizability of SSL-based models to a vasculature dataset consisting of different intensity distributions, and demonstrate that the SSL models are more robust to dataset shifts than FSL models.

## Methods

### Model Architecture

The 3D UNet (Goubran et al., 2020; Ntiri et al., 2021; Ronneberger et al., 2015) was selected as our baseline model for all experiments in this work. Each UNet had a depth of 4, with 64, 128, 256, and 512 filters at each layer. During SSL pretraining, the encoder of the 3D UNet was connected to a classifier, prior to pretraining on an auxiliary task (**Fig. 1**). For the reconstruction task, the encoder was passed to a 3D UNet decoder. After pretraining, the weights of the pretrained encoder were passed to a UNet with a randomly initialized decoder.

**Figure 1.**
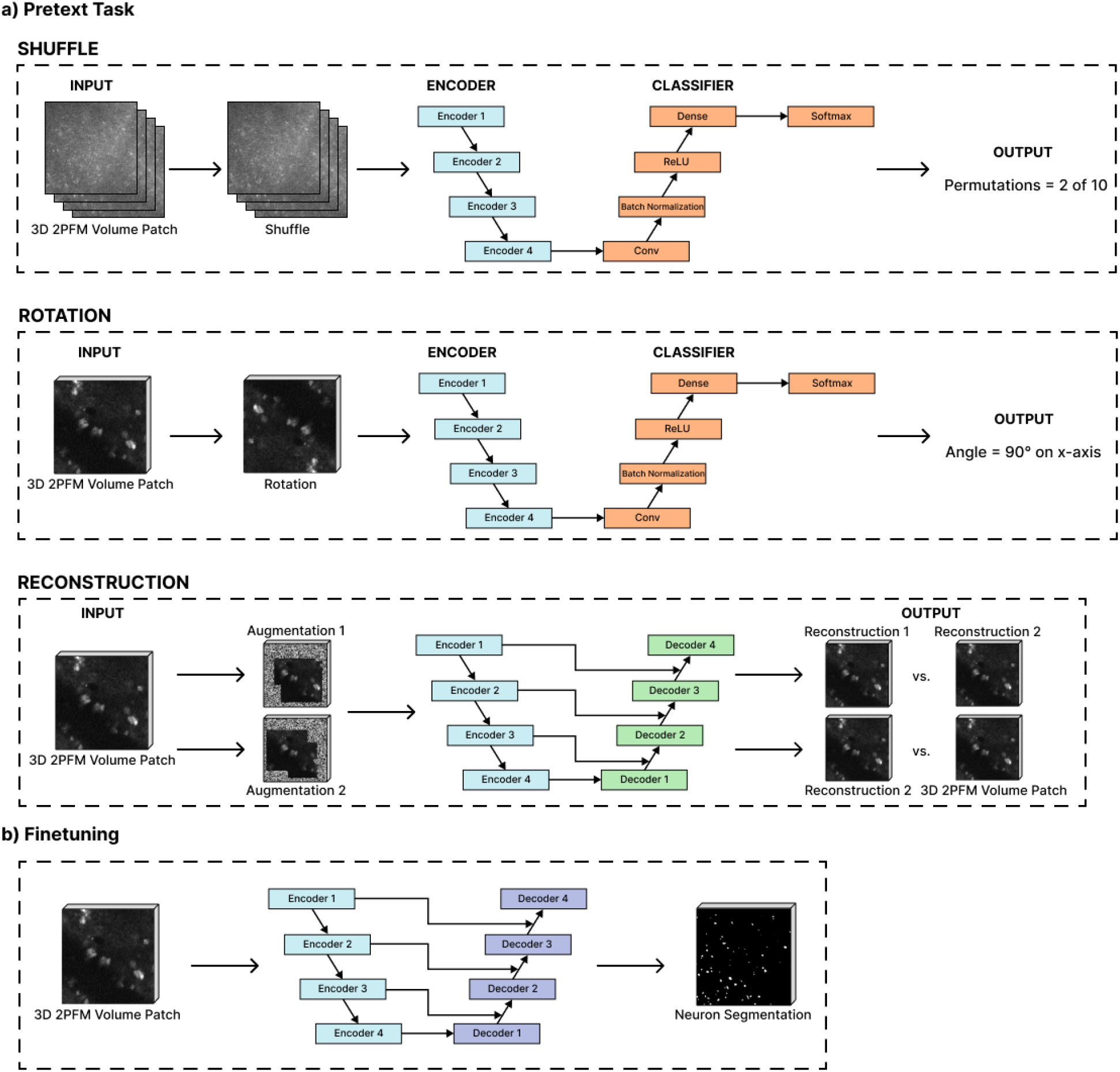
Methodology overview for training (a) and testing (b) SSL models on TPFM data. a) Segmentation models were first trained to learn a pretext task without supervision. b) The weights of pretrained models were then transferred to a segmentation model, which was trained using label supervision.

### Imaging & Datasets

We focused on two applications: segmentation of neuronal cell bodies and vessels in TPFM. The first dataset consisted of 42 volumes acquired from 25 Thy1-ChR2-YFP mice (#007612, line 18, Jackson Library) (Arenkiel et al., 2007). Mice were anesthetized with isoflurane, and implanted with a cranial window over the forelimb region in the primary somatosensory cortex, as described previously in (Rozak et al., 2024). Texas Red dextran (70 kDa MW, Thermo Fisher Scientific Inc, Waltham MA) was diluted in PBS and injected intravascularly at 33 mg/kg via a tail vein catheter to label the blood plasma. Mice were imaged on FVMPE-RS microscope (Olympus, Japan) using a 25x/1.05NA objective. An Insight DS+ tunable Ti:Sapphire laser (Spectra Physics MKS instruments Inc, USA) was used to excite both Texas Red-labeled vasculature and YFP-labeled ChR2-expressing pyramidal neurons at 900 nm. Two visible light continuous wave sapphire stimulation lasers were used to excite ChR2 at either 458 nm or 552 nm (Coherent, USA). A stack of slices was acquired from the cortical surface to a depth of 250 μm, and from 250μm to 500 μm. Images that were acquired at the cortical surface contained the larger vessels of the pial layer, with cortical penetrating arterioles descending into the parenchyma (Fumagalli et al., 2014). Channelrhodopsin-2 was predominantly expressed by pyramidal neurons in cortical layer 5 and 2/3 (Mester et al., 2019; Rozak et al., 2024).

Each image in the dataset had two channels: a channel labeling blood vessels, and another labeling neurons (**Fig.2**). Each volume had a nominal spatial resolution of 0.99 μm in plane and 2.64 μm through plane (orthogonal to the cortical surface). Ninety-six slices were acquired via resonant scanning with 5x frame averaging in plane and were later upsampled to 254 pixels through plane to achieve an isotropic image. To generate the segmentation masks for each channel, images were annotated using Ilastik (Berg et al., n.d.), a software tool that combines manual annotations with a random forest classifier. Annotations took approximately two hours per volume to complete by an experienced rater (M.R.). Each image in the dataset spanned a volume of 512x512x250 μm.

**Figure 2.**
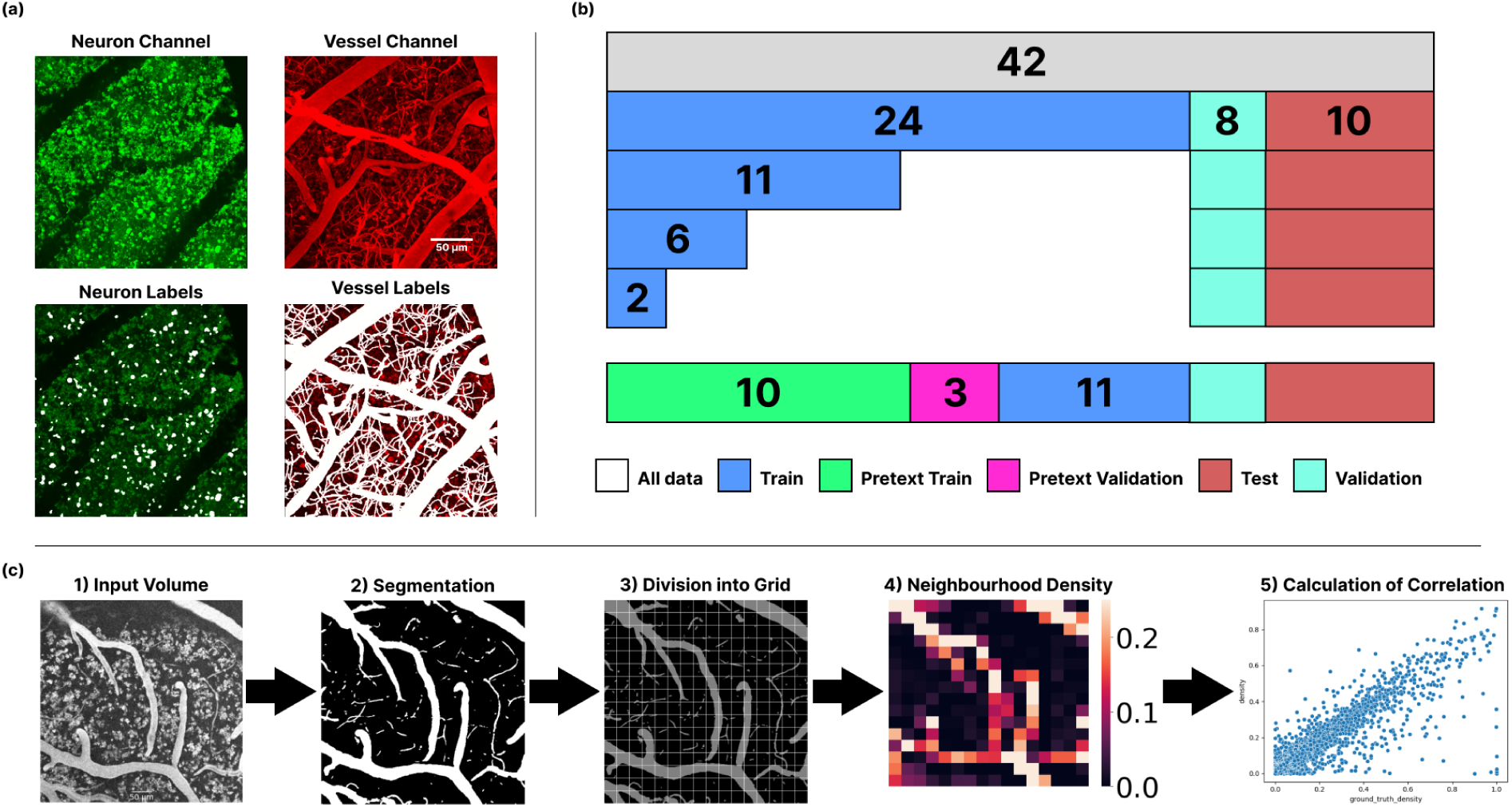
Imaging dataset and distribution for model training: a) Example maximum intensity projections of images from a Thy1-YFP mouse dataset acquired in S1FL. From left to right: (top row) image, neuron channel, vessel channel, (bottom row) ground truth labels from neuron channel, ground truth labels of vessel channel. b) Layout of data distributions used for comparison. The dataset (in grey) was split into training (blue), validation (light blue), and test (red) datasets. To train the pretext model, a portion of the training dataset was used for pretext task training (green) and validation (pink). The pretraining data was not used during finetuning of the SSL models. c) Density pooling evaluation metric. Given an input volume (1), DL models generated segmentation outputs (masks) (2). (3) The segmentation masks were then divided into neighbourhoods using a grid defined based on the size of the structure of interest, and (4) the density was calculated within each neighborhood. (5) The correlation between densities of ground truth and segmentation masks was finally computed at corresponding neighbourhoods.

The volumes in the dataset were split into training (70%), validation (10%), and test (20%) datasets (**Fig, 2**b). Thirteen (55%) volumes were removed from the training dataset for SSL pretraining. The 11 (∼45%) remaining volumes were used for both finetuning of SSL models and FSL model training. To assess the performance of the pretrained weights using different dataset sizes, the 11 volumes were reduced further to 5%, 10%, and 25% of the dataset (1, 2, and 5 volumes, respectively). The 13 SSL pretraining volumes were added to the 11 volumes to train the FSL models with 100% of the training data. Eighteen volumes were set aside for validation and testing during the finetuning. As multiple volumes were acquired from each mouse, volumes were grouped by animal prior to splitting amongst training, validation, and test datasets.

We were also interested in the effect of including data with artifacts in the training dataset. Random distributions of training data were selected from the data previously allocated for finetuning the pretrained models. The distributions had similar amounts of data as the aforementioned training datasets (i.e. 10% and 25% of the training dataset), but contained images with imaging artifacts, such as bleedthrough.

The second, out-of-sample test set used was the MiniVess dataset (Poon et al., 2023). The MiniVess dataset is an open-access annotated dataset of the cerebrovasculature of two rodent species that were acquired using TPFM. It consists of 70 3D TPFM volumes, with corresponding segmented ground truth. The volumes were acquired from the C57BL/6 and CD1 strains (20-30 g) of adult mice (N=63), and EGFP Wistar rats (Wistar-TgN(CAG-GFP)184ys) (310-630 g) (N=7)(Hakamata et al., 2001). Ground truth segmentations of the vasculature were created using thresholding methods, a 2D UNet, and manual edits.

### Preprocessing of image data

As the first dataset consists of two channels, respective channels were used for the object of interest (i.e. the channel labelling the blood vessels was used for vessel segmentation, and the channel showing the neurons was used for neuron segmentation).

Preprocessing steps varied between data used for pretraining and for finetuning. Pretraining data was first upsampled to achieve an image that was close to isotropic. Images were then normalized, with the highest and lowest values clipped between the 0.05th and 99.95th percentiles, and the resulting intensities scaled between zero and one. Foreground cropping was applied to exclude all boundaries that had nonpositive values. Then, for each image in the pretraining training dataset, 64 patches were extracted, with the transformations relevant to the pretext task applied (e.g. for the rotation task, each patch was randomly rotated one of 10 ways). Patches of size 96x96x96 voxels (FOV of 96x96x96 um^3) were extracted for all pretext tasks. Encoding for each transformation was stored as one-hot encoding was used for all pretext tasks other than the reconstruction task.

When preprocessing finetuning data, images were first upsampled along the z axis to achieve an image close to isotropic. Images were then normalized similar to pretraining data. We then applied one of the following transformations, selected at random: Contrast adjustment, Gaussian sharpening, Gaussian smoothing, Gaussian noise, and Histogram Shifting. Images were then flipped randomly along each of the three axes, with a 10% probability of flipping along each axis.

### Pretext tasks

#### Shuffling task

The shuffling task was based on the work done by Klinghoffer et al. (Klinghoffer et al., 2020b) (Figure 1a, see “Shuffle”), where the authors sought to segment axons that were imaged using light sheet fluorescence microscopy after using the SHIELD tissue clearing technique (Park et al., 2018). To do so, they presented an SSL task, here termed “shuffling”, where a random patch of size 64x64x8 was extracted from an input volume and “shuffled” along the z-axis. The model was tasked with classifying the order of the slices. The choice of this task was motivated by the tubular structure of the axons, which does not have a strict direction when imaged. Because there are 8!=40320 possible combinations, the task was made simpler by reducing the number of combinations to a random subset of 10. As reflected in the work by Noroozi and Favaro (Noroozi and Favaro, 2016), we enforced a minimum Hamming distance between the selected subset to ensure that no permutations were similar. In our experiments, random volumes of size 96x96x96 were extracted and shuffled along the z-axis. During training, each shuffled volume was labeled with one-hot encoding (resulting in a 10-class classification task).

#### Rotation and Axis Rotation tasks

Models trained on the rotation task were tasked with determining the angle of rotation. The rotation task was first introduced by Gidaris et al., where the task was implemented on 2D images (Gidaris et al., 2018). In the task, an input image was rotated in one of 4 possible 90 degree intervals. Models were tasked with classifying the angle of rotation. By learning the spatial representations, the models were expected to learn semantic features relevant to downstream tasks. The same approach has been applied to 3D datasets (Taleb et al., 2020). In the context of microscopic images, the characteristics of the images can enable the learning of features important for segmenting neurons and vasculature.

In this rotation task, a patch was randomly extracted from the dataset, and rotated at random in one of 10 possible ways. Each of the possible rotations was a multiple of 90 degrees along the x, y, and z axes, along with the origin position. In this task, we aimed to minimize the cross entropy loss. As an extension of the rotation task, we introduced another task, called axis rotation so as to test if the model could learn visual representations by predicting the axis along which image volumes are rotated. We extracted random volumes of size 96x96x96 from the training dataset, and rotated the patches at a random multiple of 90 degrees along the x, y or z axis. The model was tasked with determining the axis of rotation.

#### Reconstruction

In the reconstruction, or inpainting task, models were trained to fill in noisy, or incomplete content of an image. This task was motivated by the hypothesis that models that can successfully perform this task would be able to understand the relevant context of visible regions in the image and apply it to fill in missing areas. Pathak et al. (2016) demonstrated that a context encoder trained on the inpainting task on 2D images can learn a meaningful representation of the missing content.

The reconstruction task used in this study was taken from (Tang et al., 2022). Given a 3D volume, two sets of augmentations were applied to the same random 3D volume, generating two images. The model was tasked with generating two reconstructions of the original 3D image, while ensuring the reconstructions were as close to each other as possible. The augmentations used were in-painting, out-painting, and noise augmentation by local pixel shuffling.

### Implementation Details

All models and pipelines were developed as implemented in PyTorch and MONAI. Training and optimization were performed on the Narval cluster of Calcul Quebec and the Digital Research Alliance of Canada, with each node using 498 GB of RAM and 1 Nvidia A100 GPU, with 40GB HBM2 VRAM. An Adam optimizer was used, with different learning rates for both pretraining and finetuning (Suppl Table 1). Hyperparameters were determined using a grid search. Models were trained for 1000 epochs, with early stopping after 100 epochs without improved validation loss. For the pretext tasks, the networks were trained using Cross Entropy loss. During finetuning, 4 subvolumes of size 96x96x96 were extracted at random from each training volume. For downstream tasks, a combination of Dice and Cross Entropy loss was used. The downstream learning rate was set to 1e-4.

### Evaluation Metrics

To quantitatively evaluate the performance of all models, we used the Dice score, Precision, Recall, and Hausdorff distance (HD95), all of which are commonly used evaluation metrics in segmentation. The metrics are defined as follows:

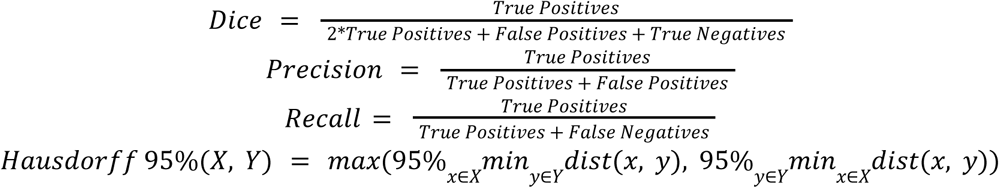

In addition, we explored more methods for quantitatively measuring the accuracy of the segmentation masks. This was done in consideration of the limitations of traditional metrics in assessing the feasibility of segmentations in downstream studies. Given these limitations, we propose an additional biologically-relevant metric for each of the segmentation tasks, namely “neighborhood density”.

To generate the neighborhood density metric, each segmentation that was generated by the models was divided into non overlapping neighborhoods of different sizes, based on the object that was being segmented (Fig 2.c). This was done to evaluate if smaller objects, such as neurons, were segmented well, while not being assessed at the same scale as vessels, which take up more space in the volumes. The density of foreground voxels within each of the neighborhoods was calculated by dividing the total number of foreground voxels by the size of the neighborhood. We then computed the Pearson correlation coefficient between the densities of the predicted segmentations and the ground truth labels. A coefficient closer to 1 translates to the model’s neighborhood density being closer to the ground truth, while a coefficient close to 0 would indicate general disagreement between the model’s segmentation and the ground truth.

## Results

We first demonstrated that our pretrained models learned features relevant to downstream tasks, while exploring the relationship between pretraining performance and finetuning. We tested our methods on segmentation of both neurons and vessels on an internal dataset. We then tested the generalizability of all trained models on an OOD dataset of mice and rat volumes (Poon et al., 2023). We used different random seeds to explore model stability and the statistical significance of SSL improvements. We further explored how well the pretext models perform when finetuned with a noisy dataset with low CNR. Finally, we tested the models trained with low CNR on the OOD dataset, assessing the robustness of our SSL models to pretraining data quality.

### Pretext Task Performance

We first assessed the accuracy of the pretext task models on their respective validation datasets. t-SNE analysis was performed to observe the layer embedding representations of the UNet encoder. The pretext model trained on the Shuffle task was better able to distinguish between classes than the models trained on the Rotation and Axis Rotation tasks (Suppl Fig 1a). Models pretrained on the Rotation and Axis Rotation tasks were also able to distinguish between classes, albeit with more difficulty. The model pretrained on the Rotation task generally failed to distinguish between classes that were similar in appearance, such those representing object rotation along the x-axis. Examining the accuracy as a function of the number of epochs (Suppl Fig 1b), the model trained on the Shuffle task was able to distinguish between classes in less than 50 epochs, reflecting the simplicity of the task relative to the Rotation and Axis Rotation tasks. Although accuracy couldn’t be used to compare the reconstruction task, the declining reconstructive loss training curve indicates that the model successfully reconstructed corrupted TPFM volumes (Suppl Fig 2).

### Neuron Segmentation Within Distribution

We finetuned and tested our pretrained models on the neuron channels in the TBI dataset. We report the Dice, precision, recall, and HD95 for each of the methods, for all subsets of finetuning training data in Suppl Table 2. The Kruskal-Wallis H-test was applied to assess statistically significant differences between the SSL models trained using 25% and 50% of the training dataset, and the FSL model trained using 100% of the dataset.

The model pretrained on the Reconstruction task with 50% of the data had a better performance than the FSL models that were trained on 100% and 50% of the dataset (Figure 3a).

**Figure 3.**
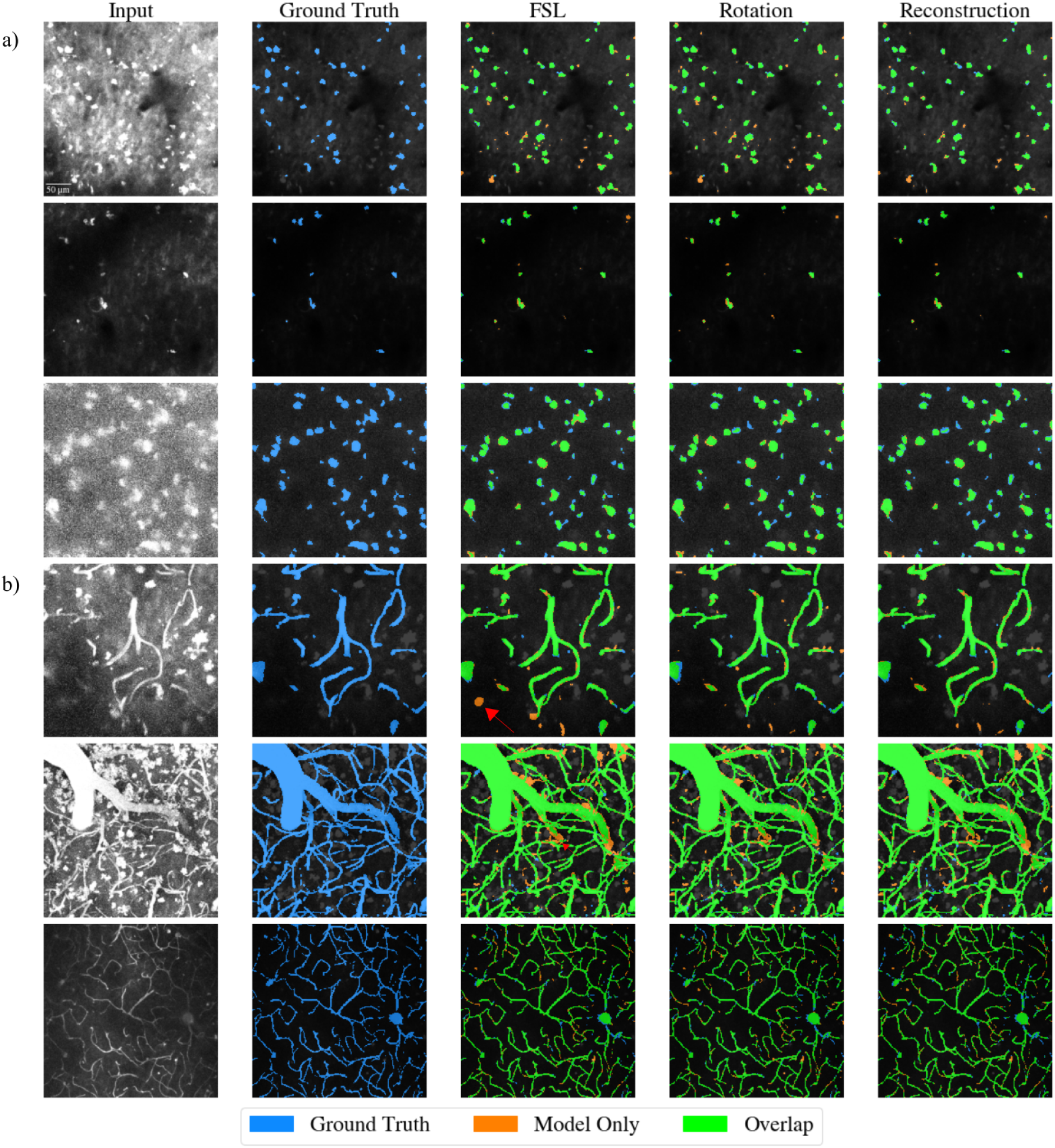
Example (a) neuron and (b) vessel segmentation masks generated by the FSL model and the SSL models pretrained on both the Rotation and Reconstruction pretext tasks and finetuned with 50% of the training dataset. From left to right: Input image, ground truth labeled mask overlaid on input image, segmentation mask generated by FSL model overlaid on both the ground truth and input image, segmentation mask generated by rotation model overlaid on both the ground truth and input image. Ground truth is blue in all images. Segmentation overlap between ground truth and segmentation maps is labeled as green. Areas segmented only by models are shown in orange. The FSL model tended to over-segment when tested on both neuron and vessel inputs (seen in red arrows).

Specifically, the Reconstruction model had a significantly higher Dice score (0.816 ± 0.044) vs. the FSL model trained with 100% of data (X+-Y) (p < .01), and lower HD95 (4.425 ± 5.118) vs FSL (X+-Y) (p < .01) (**Fig. 4**a). While the Reconstruction models did not have a significantly higher precision than FSL models, they achieved better recall (Δ_improvement_ = 0.044).

**Figure 4.**
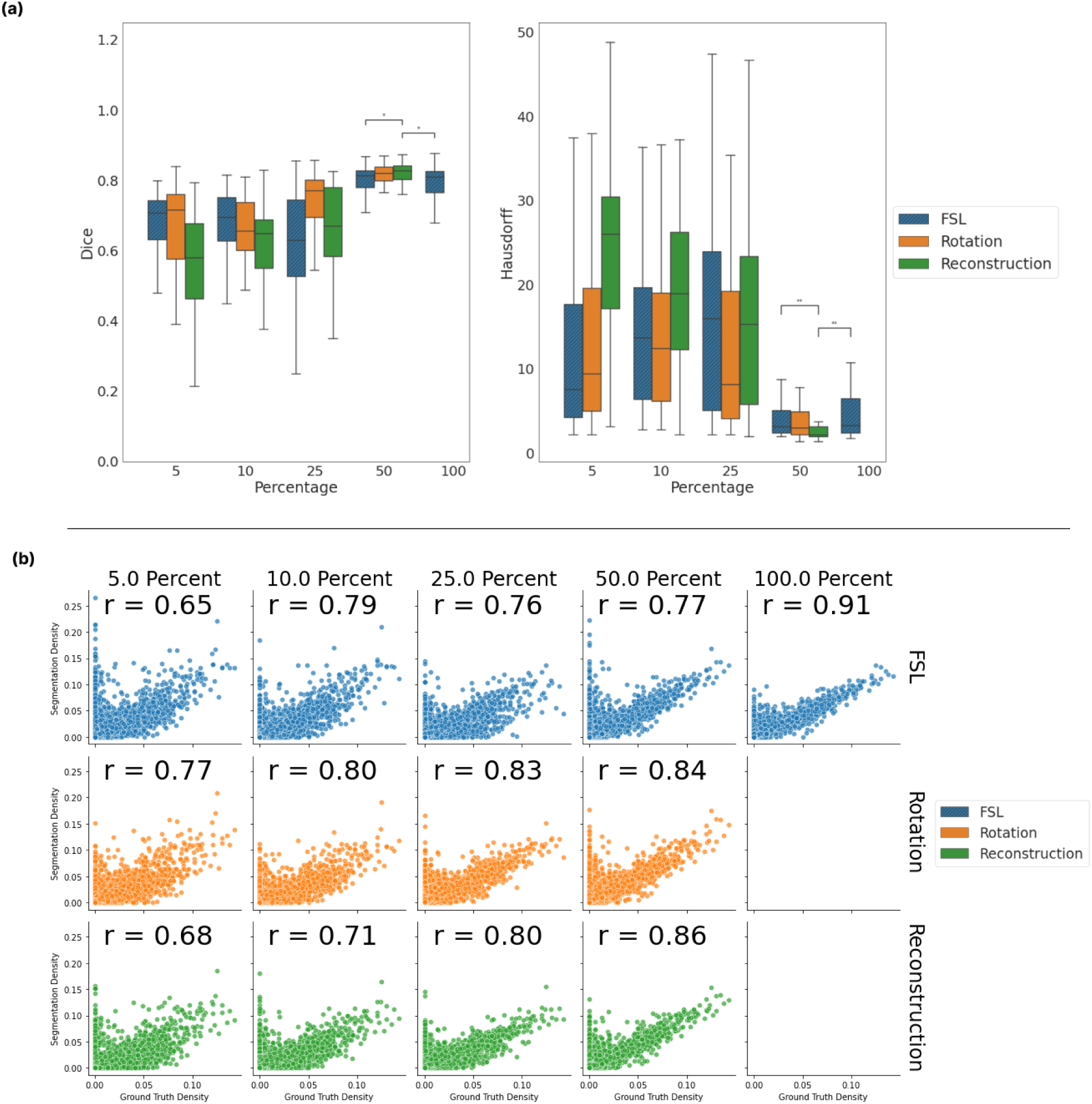
Comparison of neuron segmentation models reveals better performance of SSL models. a): Boxplot of Dice coefficient and Hausdorff Distances between ground truth neuron labels and segmentation masks generated by the FSL models, Rotation-based SSL models, and Reconstruction-based SSL models. Segmentations of 588 unique patches with shape 96^3 voxels were used. All differences in scores with a *p-value* < 0.01 were considered significant. b): Scatterplot of segmentation neighborhood densities plotted against ground truth density. Subvolumes were set to (32, 32, 32). Each row is a different method of training the model; each column is a percentage of training data used during finetuning. Inline: the Pearson R correlation coefficients between the pooled segmentations and the pooled ground truth.

The neighborhood density results revealed a high degree of agreement between the labels generated by the SSL models and the ground truth (Fig. 6). The Rotation and Reconstruction models that were finetuned on 25% and 50% of the training dataset had a higher correlation with the ground truth densities (Rotation - 25%: 0.83; 50%: 0.84, Reconstruction - 25%: 0.80, 50%: 0.86) than did the FSL models trained with the same amounts of training data (25%: 0.76, 50%: 0.77), suggesting that the SSL-pretrained models generate label maps that are more beneficial for downstream analysis tasks such as investigating alterations in neuronal densities.

### Vasculature Segmentation Within Distribution

All evaluation metrics and corresponding box plots for models trained on the vessel channel are summarized in Table 1, and Figure 5.a. All SSL models finetuned on 50% of the training dataset had comparable Dice coefficients to the FSL model trained on the same amount of data (no significant differences, p > 0.05). Of the pretrained models, the model pretrained on the Rotation task achieved the highest Dice score (0.878 ± 0.088). Both the Rotation and Axis Rotation pretrained models had comparable Dice scores to the FSL models with increasing amounts of finetuning data. The neighborhood density maps revealed a high degree of similarity between the SSL models and the ground truth (**Fig.5**.b), suggesting that the segmentation maps generated by the SSL models were on par with those produced by the FSL models.

**Figure 5.**
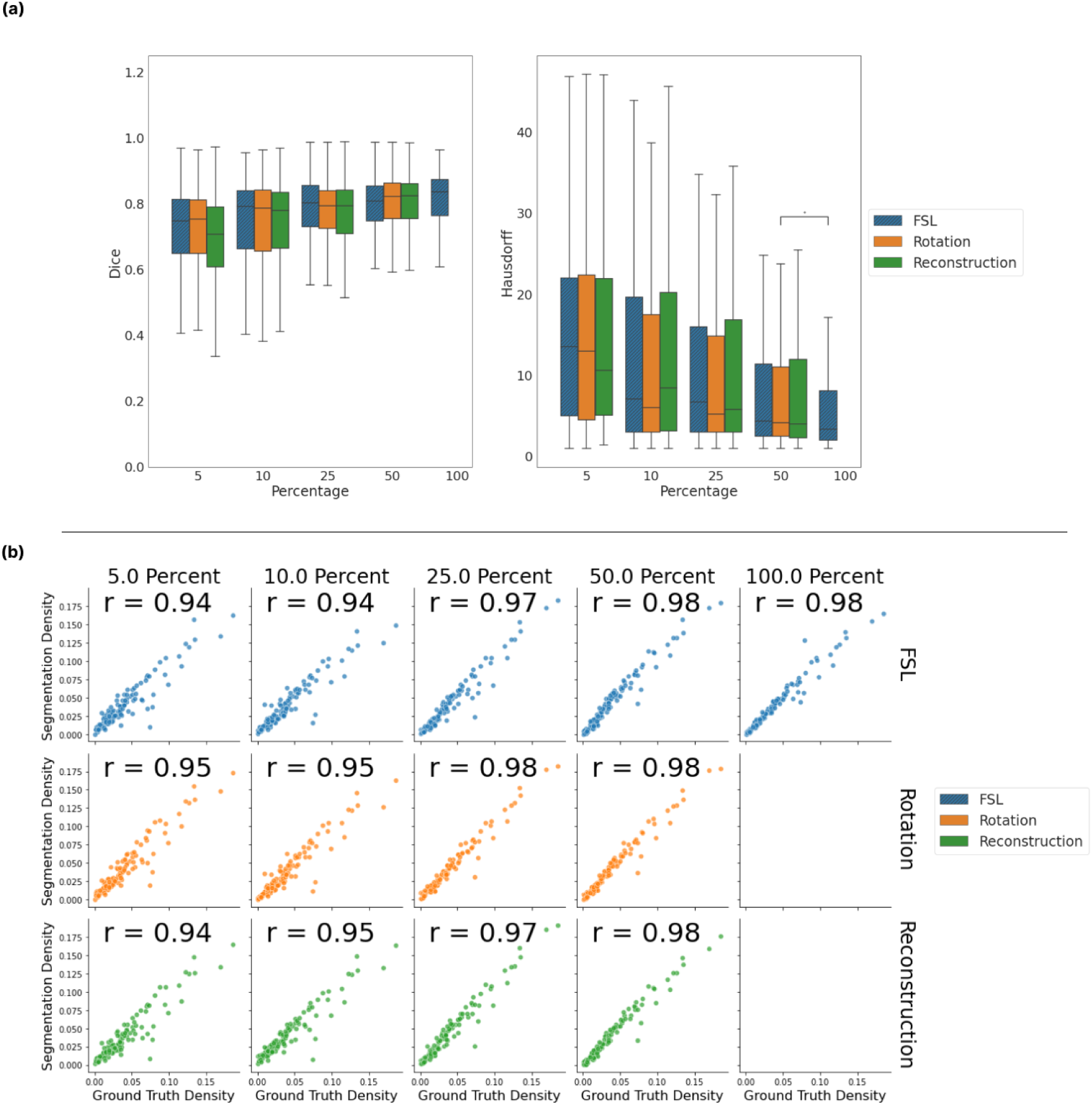
Comparison of vessel segmentation models reveals similar performance between SSL and FSL models. a) Boxplot of Dice coefficient and Hausdorff Distances between ground truth vessel labels and segmentation masks generated by the FSL models and the SSL models pretrained on the Rotation and Reconstruction tasks. Segmentations of 718 unique patches with shape 96^3 voxels were used. b) Scatterplot of segmentation neighborhood densities plotted against ground truth density. Subvolumes were set to (128, 128, 128). Each row is a different method of training the model, each column is a percentage of training data used during finetuning. Inline: the Pearson R correlation coefficients between the pooled segmentations and the pooled ground truth.

**Table 1.**
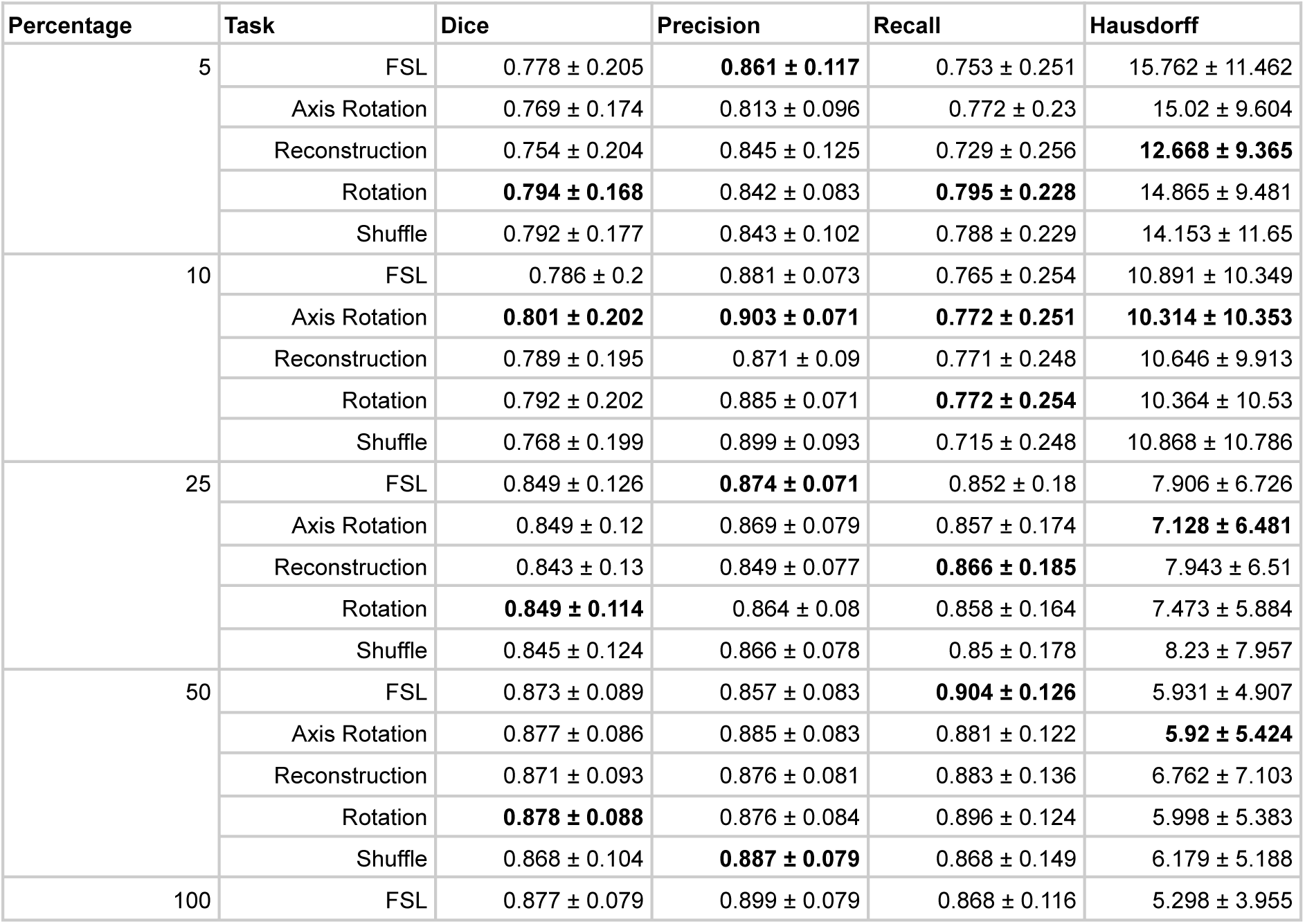
Mean evaluation metrics across different models on vessel segmentation in the test dataset.

### Effects of Initialization

We explored the stability of our pretext models to ensure that our results were not influenced by random model initialization. All pretrained models were finetuned with three different random seeds with the aforementioned proportions of data. The models were then tested on the test dataset. Boxplots of the results did not reveal large deviations between the different models produced from different seed initializations (Supplementary Figures 10 and 11). For models trained on either the neuron or vessel segmentation task, the Kruskal Wallis test did not reveal any differences between the runs (p > 0.05), indicating the stability of SSL models in relation to model initialization.

### Effects of Data Quality on Model Performance

In this experiment, we investigated the relationship between model performance and dataset CNR. We selected visually noisy subsets with a large amount of crosstalk, and set them in groups that matched the original datasets of size 10%, 25%, and 50% (**Fig. 6**). These datasets will be hereafter referred to as “noisy”, while models trained on the original datasets will be termed “original ’’. The new datasets were then used to finetune the pretrained models, with the same preprocessing steps applied.

**Figure 6.**
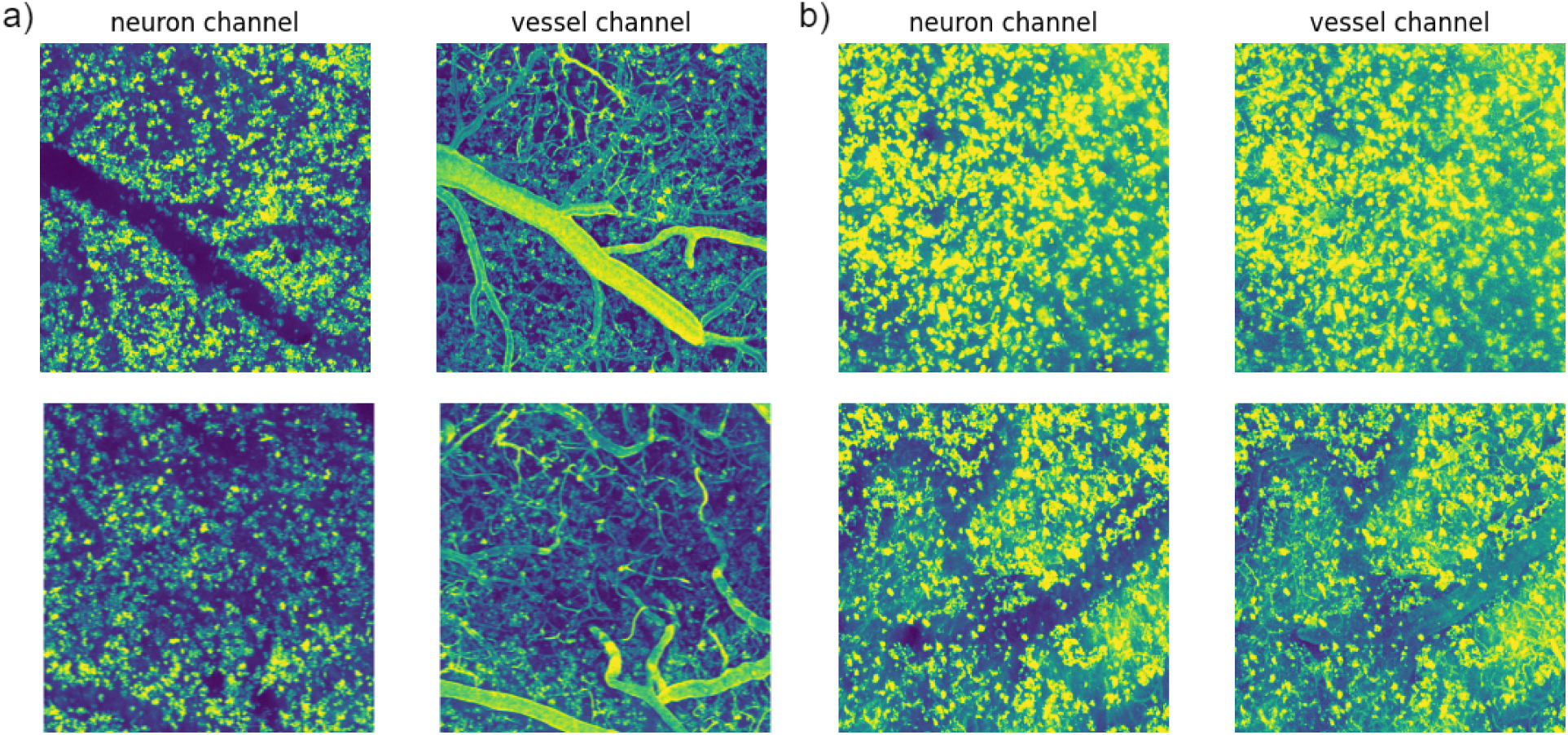
Example maximum intensity projections of images used in (a) original and (b) noisy training datasets. (a) Maximum intensity projections of (left) neuron and (right) vessel volumes used in the original training dataset. (b) Maximum intensity projections of both the neuron and vessel channel of two example volumes used in the ‘*noisy’* training dataset. A high degree of bleedthrough from the neuron channel to the vessel channel can be observed in the vessel channel (right).

When finetuned and tested on the *neuron* dataset, models trained on the noisy dataset (whether SSL or FSL) outperformed those models trained on the original data. SSL models trained on the noisy dataset outperformed the FSL models. In particular, the model pretrained on the Reconstruction task and finetuned with 50% of the noisy dataset had a Dice of (0.823 ± 0.044), compared to the FSL model trained with the same amount of noisy data (0.799 ± 0.057) (p < 0.05). The difference in Dice performance between the noisy and original datasets decreased as the amount of finetuning data increased. A tradeoff between precision and recall was observed in the FSL model finetuned with the full noisy dataset compared to the FSL model trained with the original dataset (**Fig. 7**). While the FSL model trained with 50% of the noisy dataset had a higher precision (0.92 vs 0.89), it had a lower recall (0.72 vs 0.74). The Recall of the Reconstruction-based SSL model did not decline when the model was finetuned on 50% of the noisy dataset. This was not true for either the FSL models or the other SSL models.

**Figure 7.**
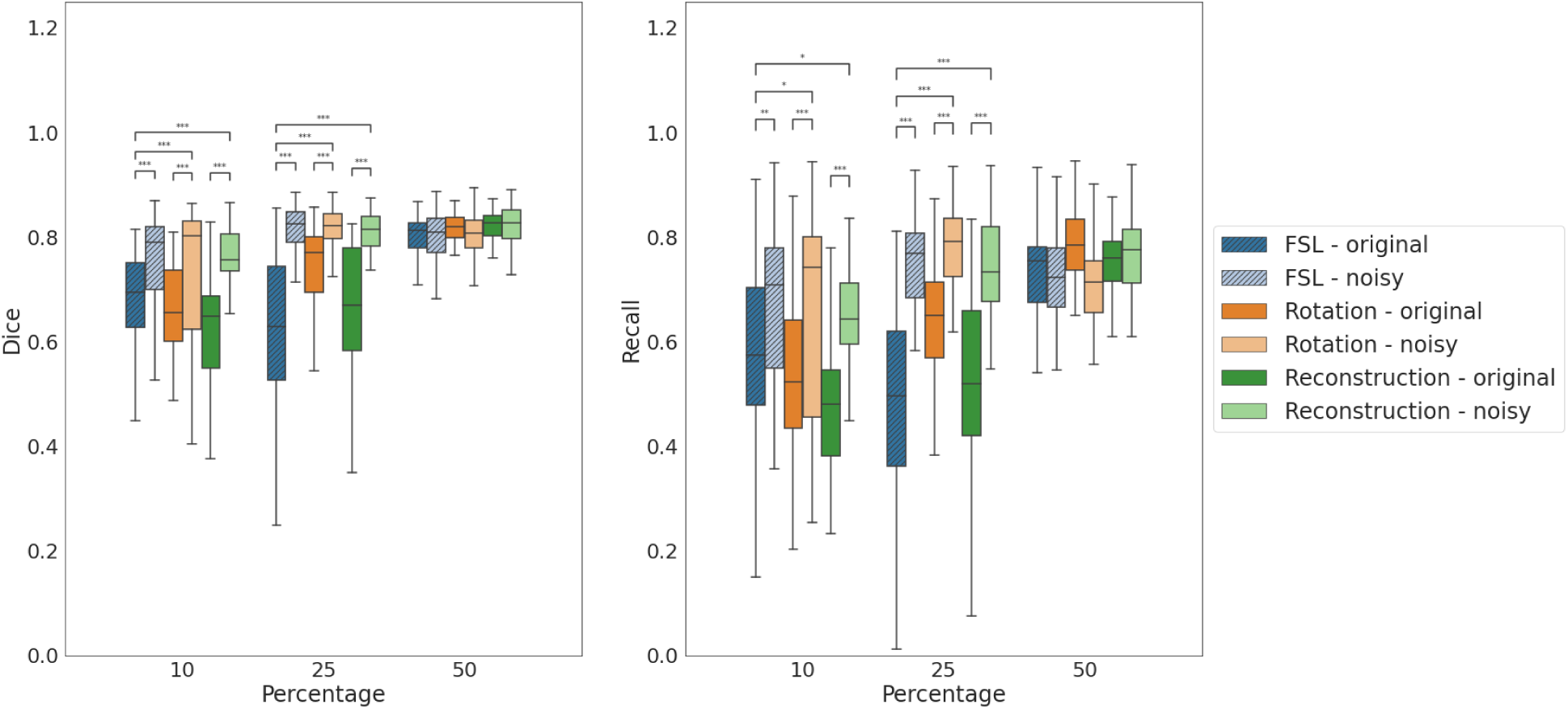
Boxplots of Dice coefficient and Recall between ground truth neuron labels and segmentation masks generated by the FSL models and SSL models pretrained on the Rotation and Reconstruction tasks. Lighter shades designate models that were finetuned with the *noisy* dataset. Segmentations of 718 unique patches with shape 96^3 voxels were used.

Pretrained SSL models that were finetuned on the noisy *vessel* datasets performed as well as their counterparts trained with the same amounts of original data. The model pretrained on the Rotation task and finetuned with 25% of noisy data had a higher Dice (0.822 ± 0.121) than the FSL model trained with the same dataset (0.787 ± 0.162), but a lower Dice than the FSL model trained with 25% of the original dataset (0.849 ± 0.126). The results suggest that the SSL models finetuned on the noisy vessel dataset are comparable to the FSL models trained with the original dataset.

### Out-of-Distribution Vasculature Segmentation

We assessed how well models pretrained on the internal dataset generalize to out-of-distribution datasets. We used the models finetuned from the previous experiment, and tested them on the MiniVess dataset. The Dice score, Precision, Recall, and Hausdorff distances were computed for all methods (**Table 2**). When finetuned on 5, 10 and 50% of the labeled dataset, the SSL tasks had higher Dice scores than FSL models by 2%, 5% and 6%, respectively. In particular, when finetuned with 50% of the training dataset, the Reconstruction model had a higher Dice score (0.783±0.119) than the FSL model (0.768 ± 0.139) when tested on the MiniVess dataset (p < 0.01).

**Table 2.** Mean evaluation metrics across different models on when segmenting MiniVess dataset.

| Percentage | Task | Dice | Precision | Recall | Hausdorff |
| --- | --- | --- | --- | --- | --- |
| 5 | FSL | $0.698 \pm 0.15$ | **$0.892 \pm 0.08$** | $0.598 \pm 0.172$ | $10.45 \pm 9.123$ |
| | Reconstruction | $0.578 \pm 0.158$ | $0.87 \pm 0.097$ | $0.453 \pm 0.163$ | $14.657 \pm 9.289$ |
| | Rotation | **$0.727 \pm 0.141$** | $0.842 \pm 0.103$ | **$0.671 \pm 0.174$** | **$8.68 \pm 8.664$** |
| 10 | FSL | $0.661 \pm 0.199$ | $0.9 \pm 0.1$ | $0.554 \pm 0.215$ | $14.08 \pm 10.984$ |
| | Reconstruction | **$0.715 \pm 0.168$** | $0.898 \pm 0.079$ | **$0.623 \pm 0.194$** | **$11.699 \pm 10.447$** |
| | Rotation | $0.64 \pm 0.195$ | **$0.903 \pm 0.099$** | $0.525 \pm 0.206$ | $15.73 \pm 11.496$ |
| 25 | FSL | **$0.765 \pm 0.135$** | **$0.898 \pm 0.084$** | **$0.691 \pm 0.167$** | **$8.224 \pm 8.236$** |
| | Reconstruction | $0.712 \pm 0.152$ | $0.868 \pm 0.101$ | $0.63 \pm 0.183$ | $10.381 \pm 7.858$ |
| | Rotation | $0.681 \pm 0.153$ | $0.891 \pm 0.092$ | $0.576 \pm 0.181$ | $12.066 \pm 8.367$ |
| 50 | FSL | $0.768 \pm 0.139$ | $0.803 \pm 0.116$ | **$0.781 \pm 0.185$** | **$8.114 \pm 9.682$** |
| | Reconstruction | **$0.783 \pm 0.119$** | **$0.887 \pm 0.091$** | $0.723 \pm 0.153$ | $8.144 \pm 8.38$ |
| | Rotation | $0.778 \pm 0.132$ | $0.869 \pm 0.099$ | $0.73 \pm 0.166$ | $8.142 \pm 8.531$ |
| 100 | FSL | $0.743 \pm 0.143$ | $0.889 \pm 0.092$ | $0.665 \pm 0.168$ | $9.013 \pm 9.482$ |

All SSL models had decreasing average HD95 distances with increasing amounts of training data. The SSL models finetuned on 5% and 10% of the labeled data had lower HD95 scores than the FSL models trained with corresponding amounts of labeled data. The Rotation and Reconstruction models that were finetuned on 25% and 50% of the training dataset had a higher correlation with the OOD ground truth densities (Rotation - 25%: 0.89, 50%: 0.89, Reconstruction - 25%: 0.90, 50%: 0.88) than FSL models trained with the same amounts of training data (25%: 0.87, 50%: 0.88) (**Fig. 8**b), suggesting that the SSL models had a closer shape agreement to the ground truth.

**Figure 8.**
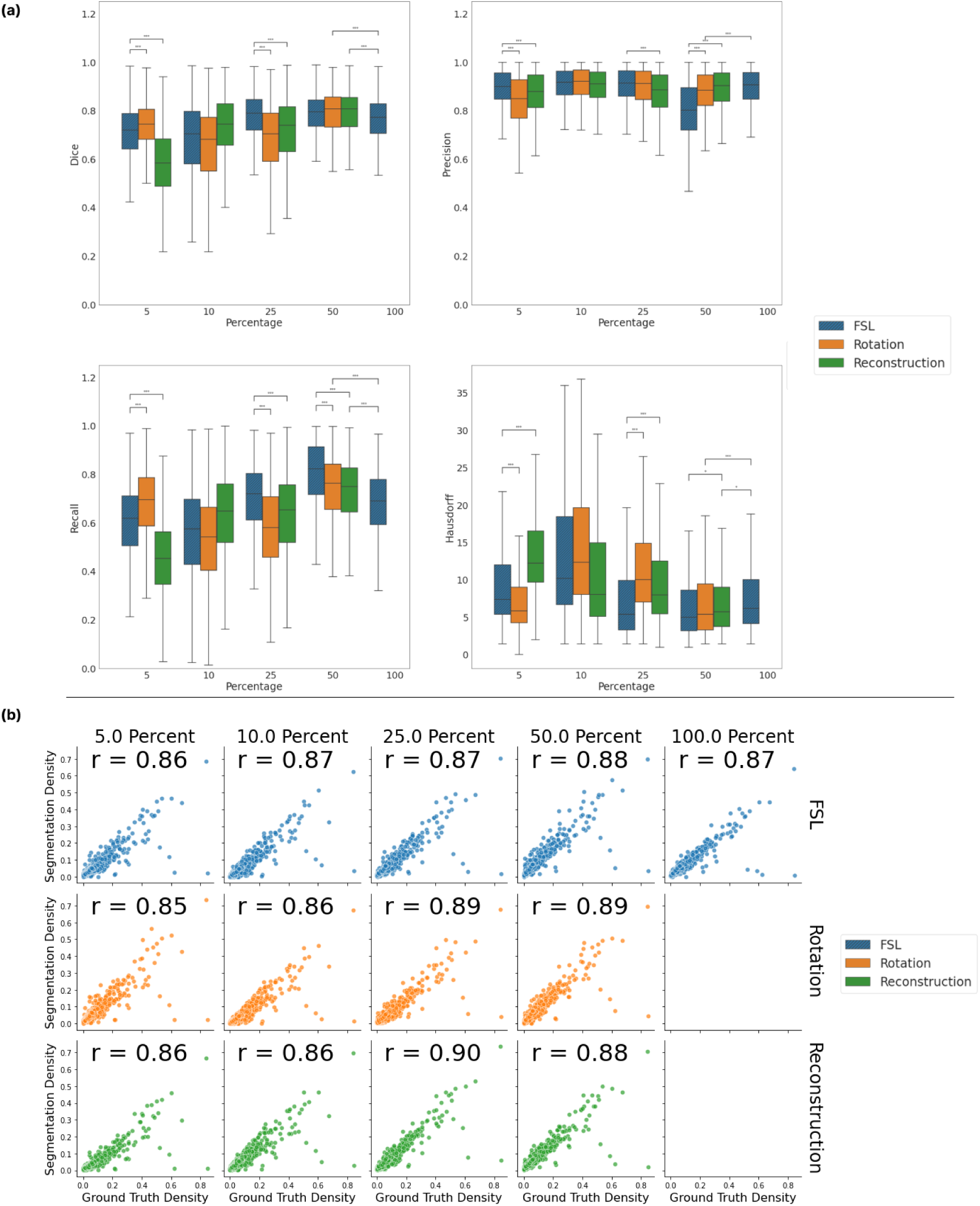
Comparison of vessel segmentation models when tested on OOD distribution. a) Boxplot of Dice coefficient, Precision, Recall, and Hausdorff Distances between ground truth vessel labels and segmentation masks generated by the FSL models and SSL models pretrained on the Rotation and Reconstruction tasks when tested on MiniVess dataset. b) Scatterplot of segmentation neighborhood densities plotted against ground truth neighborhood density from the MiniVess dataset. Volumes were divided into regions of size (128, 128, 128). Each row is a different method of training the model, each column is a percentage of training data used during finetuning. Each color represents a level of depth. Inline: the Pearson’s R correlation coefficients between the pooled segmentations and the pooled ground truth.

## Discussion

We present self-supervised models for neuronal cell body and vasculature segmentation in TPFM images. The models were first trained to perform different pretext tasks, we then employed model weights and trained them on datasets of increasing size, showing that they could segment both neurons and vessels on par or better than FSL models while using 50% of finetuning data. We found that the pretrained models were robust to degradation in training data quality compared to FSL models. We finally found that SSL models generalize better on OOD with different image characteristics than the training dataset. Pretraining DL models with SSL presents a promising avenue to achieve a robust and efficient segmentation in TPFM and applications with limited labeled data.

### SSL Task-specific Performance

In this work, we did not seek to compare our segmentation model to other SOTA models, but rather aimed to explore and demonstrate the benefits of SSL. The UNet has already served as an effective baseline model for segmenting objects of interest in 3D microscopy datasets (Poon et al., 2023; Rozak et al., 2024), due to its high performance across different tasks and ability to accurately capture local features. The features learned through the various SSL pretext tasks were demonstrated to improve when FSL models are trained to segment both neurons and vessels, in line with previous work in biomedical imaging (Chen et al., 2019; Taleb et al., 2020).

We validated the SSL pipeline in two different segmentation tasks, using two large 3D microscopy datasets. In the neuron segmentation task, the SSL model pretrained with reconstruction task outperformed the baseline FSL UNet, achieving a higher Dice score with half of the training dataset. Models pretrained on the reconstruction task learned salient information about the image volumes, despite missing a portion of relevant information due to masking. Models pretrained on the rotation task had some success distinguishing between different object orientations without context clues visible to human raters, such as the differences between slices in a given axis. SSL frameworks using the rotation task during pretraining have demonstrated success in videos and medical imaging (Jing et al., 2018; Li et al., 2021; Taleb et al., 2020), but in many of these cases, the spatial information in datasets, such as direction and shape of objects, was easier to perceive than in TPFM data.

The performance of the SSL models is likely pretext task-specific. Both the rotation task and the reconstruction task are inherently more difficult tasks than the shuffling task. The rotation task is difficult, as it requires determining both the axis, and the number of degrees of rotation. Different classifications in the rotation task are similar, such as rotating along either the Y or Z axis at 180 degrees. Interestingly, the axis rotation pretext task produced a similar performance to that of the rotation task on the vessel dataset, despite being an easier pretext task. The reconstruction task could not be compared to the other tasks using tSNE plots, as the outputs were not one hot encoded. However, the reconstruction task is more difficult than the shuffling task, as completing the task successfully requires learning both local and global features in data. We observed that pretext task difficulty, and not pretext task performance, is related to downstream model performance.

### Performance on Within and Out-of-Distribution Datasets

We found that the SSL models outperformed the FSL models on both the vessel and the neuronal segmentation tasks. We demonstrated that the model pretrained on the Reconstruction task was best at segmenting neurons in the test dataset, outperforming the FSL model on both within and OOD datasets. Models trained on either the Reconstruction or Rotation tasks performed comparably to FSL models when segmenting vessels. However, when trained on 25% of the dataset, the SSL models failed to segment vasculature as well as FSL models (p < .01). Observing the neighbourhood density maps indicated that there was a high degree of similarity between the segmentation maps of the SSL and FSL models.

Models often cannot translate performance to datasets with other distributions. This may be due to statistical shifts in the properties of the datasets, such as in intensity distributions. For example, shifts between datasets may occur from acquiring images at different step sizes, or from as a result of using aging equipment. Our results demonstrate that SSL models trained on varying amounts of data are robust to OOD datasets. The high performance of SSL models finetuned with 5% of the dataset (p < .01) demonstrates that SSL is a promising avenue to generate models that can adapt to different distributions.

The findings highlight the potential of SSL models in medical image segmentation tasks, particularly in scenarios with limited labeled data, and when tested on different distributions. Future work could focus on making the models more adaptable to different distributions at test using methods such as test time augmentation. Further investigations into the roles of data quality and diversity during pretraining may provide insights into optimizing SSL methods for broader applications.

### Effect of Noise on Generalizability

Microscopy datasets often contain a high amount of noise due to light attenuation or imaging artifacts. Images may be discarded from studies due to the inability to utilize the volumes. We thus sought to determine if images with low SNR and CNR would have some utility in training segmentation models. We found that SSL models that were finetuned on the noisy datasets had similar results to the SSL models trained on the original dataset during testing. We also observed that the models were more robust to noise.

The results show that *pretrained* models’ performance are robust against changes in training data distributions. The differences in CNR between the neuron and vessel datasets can be observed in Figure 9. As the image depth increases, the mean CNR for the test and “noisy” neuron training datasets begin to overlap, while the mean CNR for the “original” neuron training dataset remained higher. The similarity in image distributions may have translated to an improved performance. This would explain how the SSL models finetuned on noisy neuron data were able to outperform the original SSL models, with higher Dice and Recall scores (**Fig. 7**).

**Figure 9.**
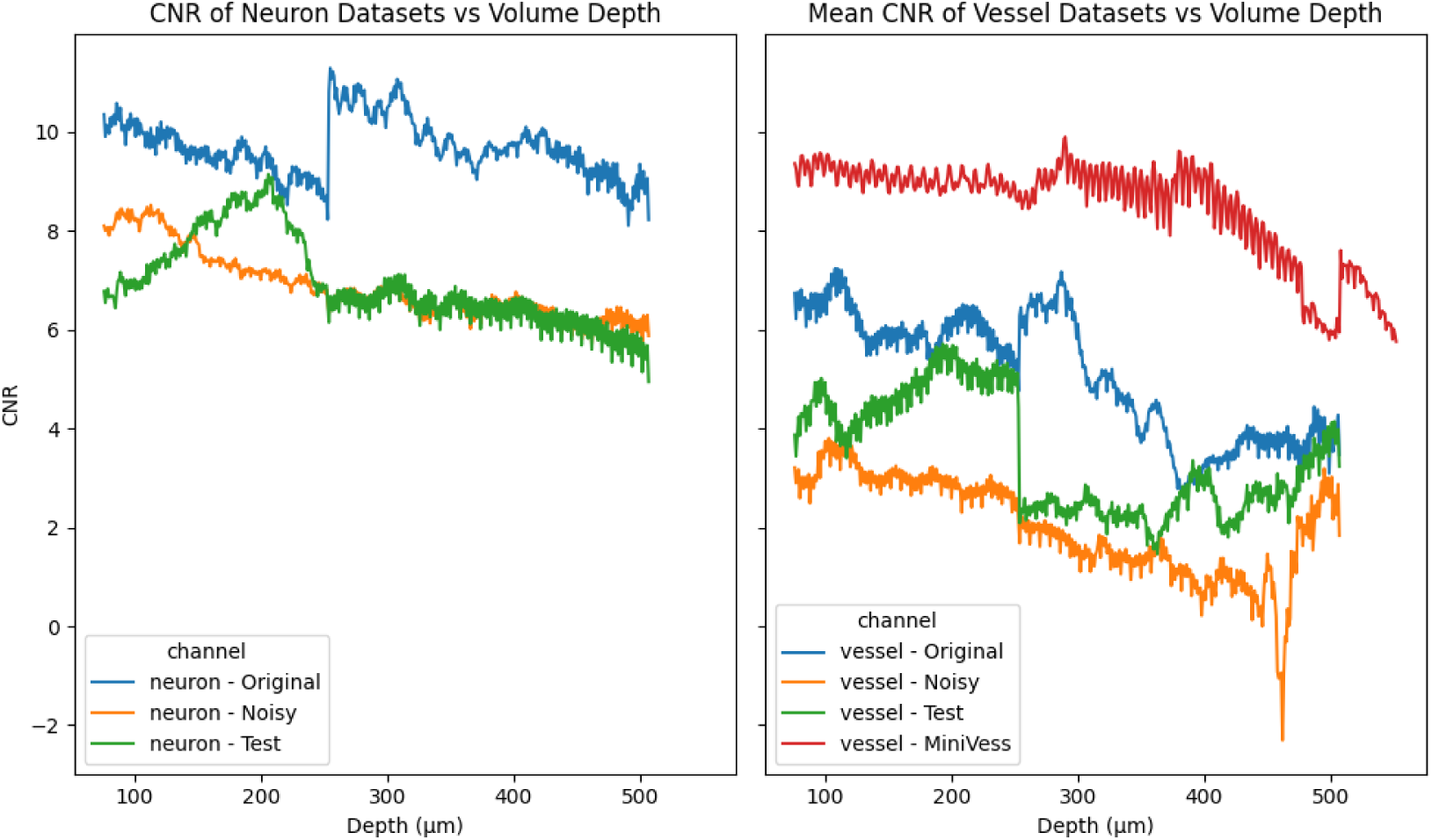
CNR distributions of the training and test datasets explain model performance on datasets. Left: Mean CNR of neuron datasets plotted as a function of volume depth reveal CNR between noisy training dataset distribution and test distribution. Right: CNR of vessel datasets plotted as a function of volume depth.

Observing the CNR of the *vessel* datasets, we found that the distributions of the original, noisy, and test datasets were similar, resulting in small differences across SSL and FSL models.

However, the SSL models that were finetuned on “noisy” data still outperformed the FSL models. These findings demonstrate that the SSL models’ performance is more robust against distribution shifts in training data. This result is important, as researchers may employ training data with a wide range of image characteristics. With relatively lower quality finetuning data, SSL models can still be finetuned to have improved performance on downstream tasks.

### Limitations

While the augmentations applied to the pretraining datasets generated large amounts of pretraining data, the size of the pretraining dataset was relatively small. Further experiments would benefit from pretraining the model on a larger volume of unlabeled data. While the metrics proposed to compare the segmentation masks and the images are promising, the neighborhood density is not specific to the object of interest, meaning that the patched density may not reflect how well a mask represents the foreground in an image. Future work will investigate other metrics using additional data representations like graphs (Damseh et al., 2019)

## Conclusion

In this work, we proposed and assessed a self-supervised pipeline for the segmentation of neurons and vasculature in TPFM data. We presented a series of pretext tasks that generate relevant representations for segmentation in TPFM. We demonstrated that models trained using SSL performed better than did the FSL ones with less data, and surpassed FSL models on out-of-distribution datasets. Finally, we demonstrated that SSL models were more robust to variations in training data quality. Our SSL pipeline is available to the research community to enable the development of robust and generalizable segmentation models for TPFM and microscopy.

## Acknowledgments

This work was supported by funding from Canadian Institute of Health Research (CIHR) project grants PJT178059, PJT376309, and PJT156179, and a Natural Sciences and Engineering Research Council (NSERC) Discovery grant RGPIN-2021-03728. . E.N. was funded by the Department of Medical Biophysics, University of Toronto’s Excellence Award. B.S. is funded by the Canada Research Chairs program award #CRC-2018-00042. M.G. is funded by the Canada Research Chairs program award #CRC-2021-00374. We are grateful to Calcul Québec and the Digital Research Alliance of Canada (alliancecan.ca) for their allocation of compute resources that in part supported this research.

## Authorship Contribution

**Emmanuel Edward Ntiri:** conceptualization, methodology, software, validation, formal analysis, investigation, writing - original draft, writing - review & editing, visualization, project administration.

**Tony Xu:** conceptualization, methodology, validation, writing - review & editing

**Matthew Rozak:** conceptualization, resources, data curation, writing - review & editing **Ahmadreza Attarpour:** conceptualization, methodology, validation, writing - review & editing **Adrienne Dorr:** resources, data curation, writing - review & editing.

**Bojana Stefanovic:** conceptualization, writing - review & editing, supervision

**Maged Goubran:** conceptualization, methodology, formal analysis, resources, writing - original draft, writing - review & editing, supervision, project administration, funding acquisition.

## Code & Data availability

The developed algorithm is available at https://github.com/AICONSlab/self_micro, under the GNU General Public License v3.0. We have developed an easy-to-use pipeline with thorough documentation for easy access for users with limited programming knowledge.

---

**Suppl Figure 1.**
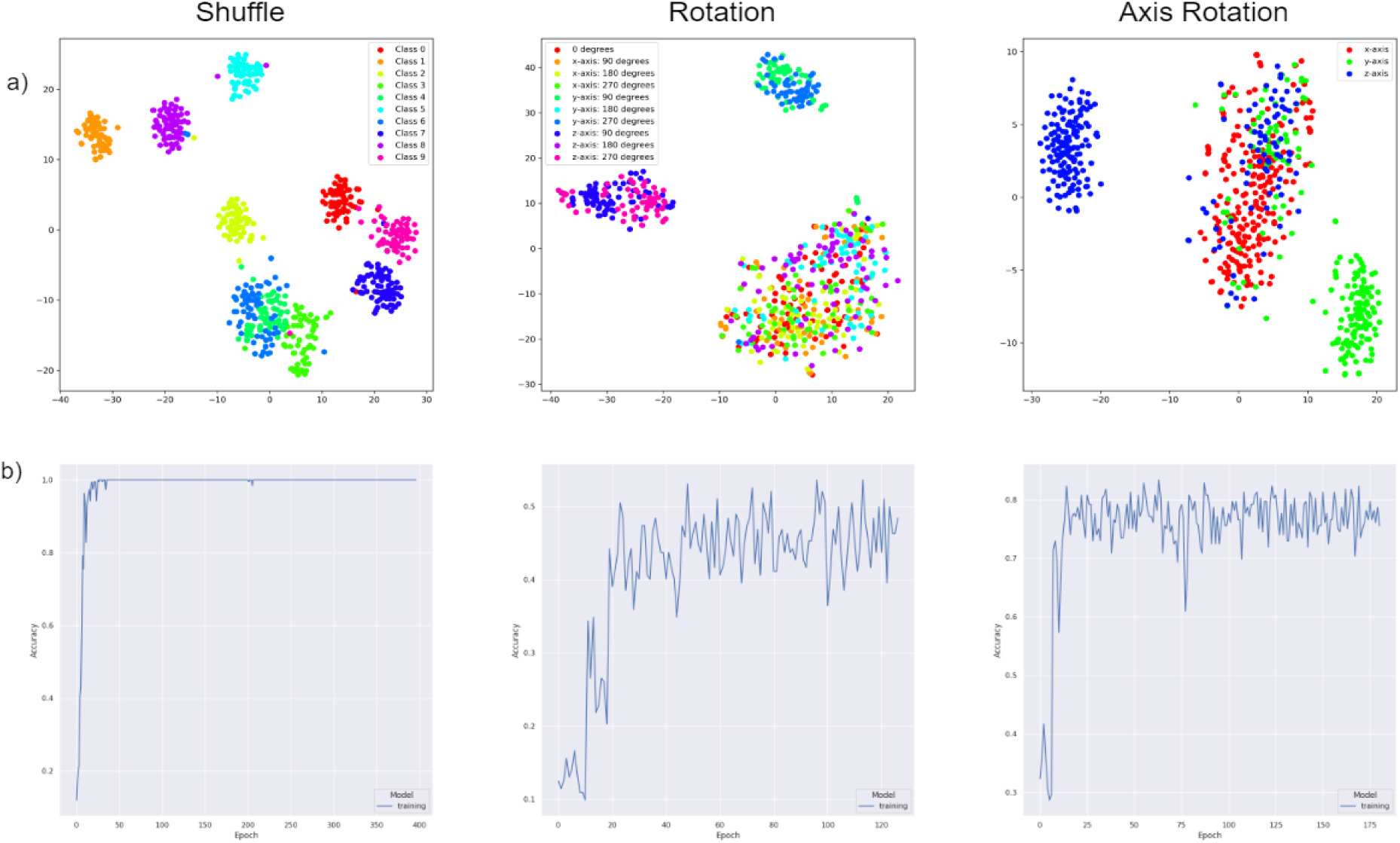
Visualization of pretraining model results. a) t-SNE plots of Shuffle, Rotation, and Axis Rotation tasks when pretrained on vessel segmentation data. Each dot in the output was coloured by a class number representing the pretext task label. b) plots with pretext task accuracy as a function of the number of training epochs used to solve the pretext task.

**Suppl Figure 2.**
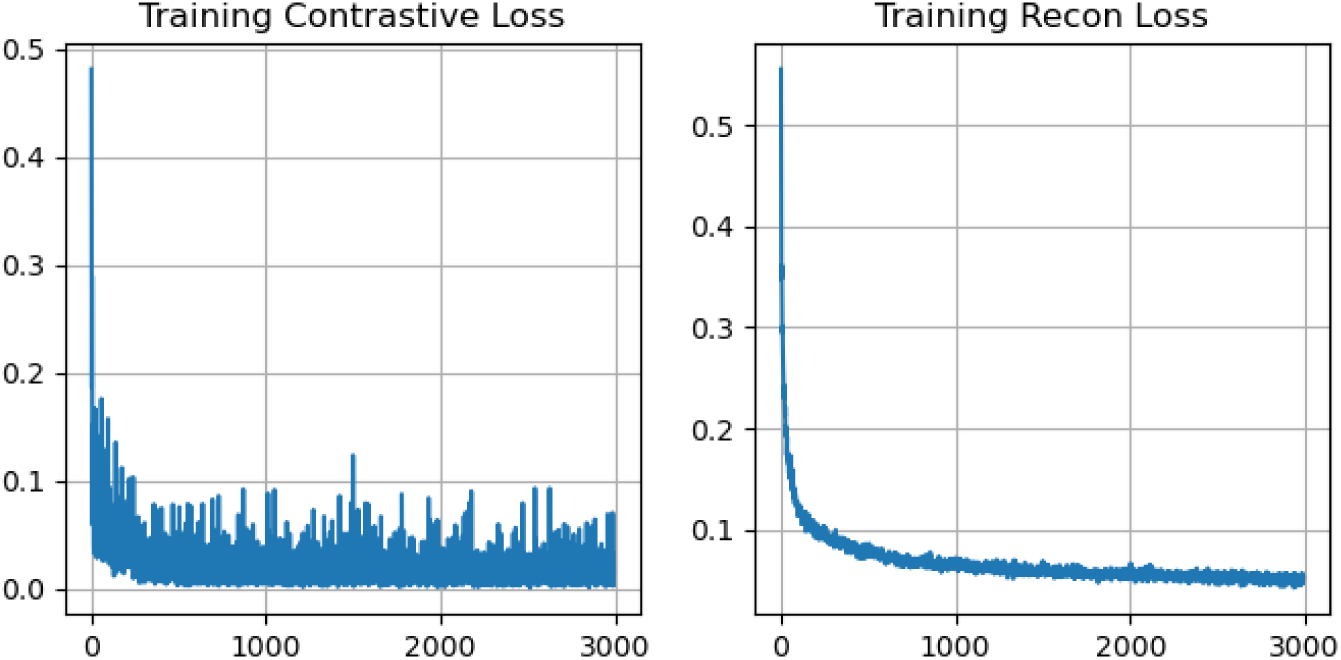
Visualization of reconstruction pretraining model results.

**Suppl Figure 3.**
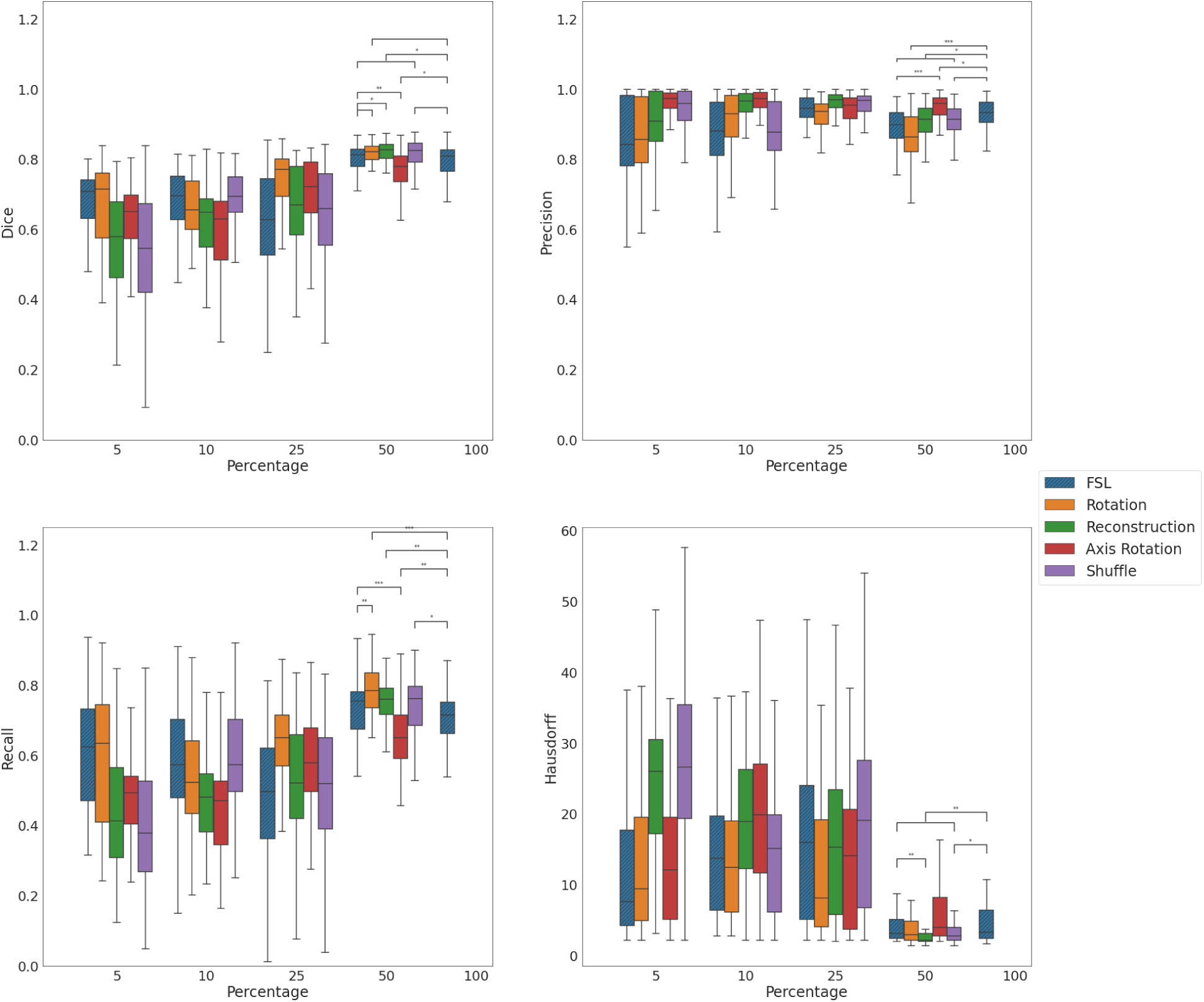
Boxplots of Dice coefficient, Precision, Recall, and Hausdorff Distances between ground truth neuron labels and segmentation masks generated by the FSL models and SSL models pretrained on all pretext tasks.

**Suppl Figure 4.**
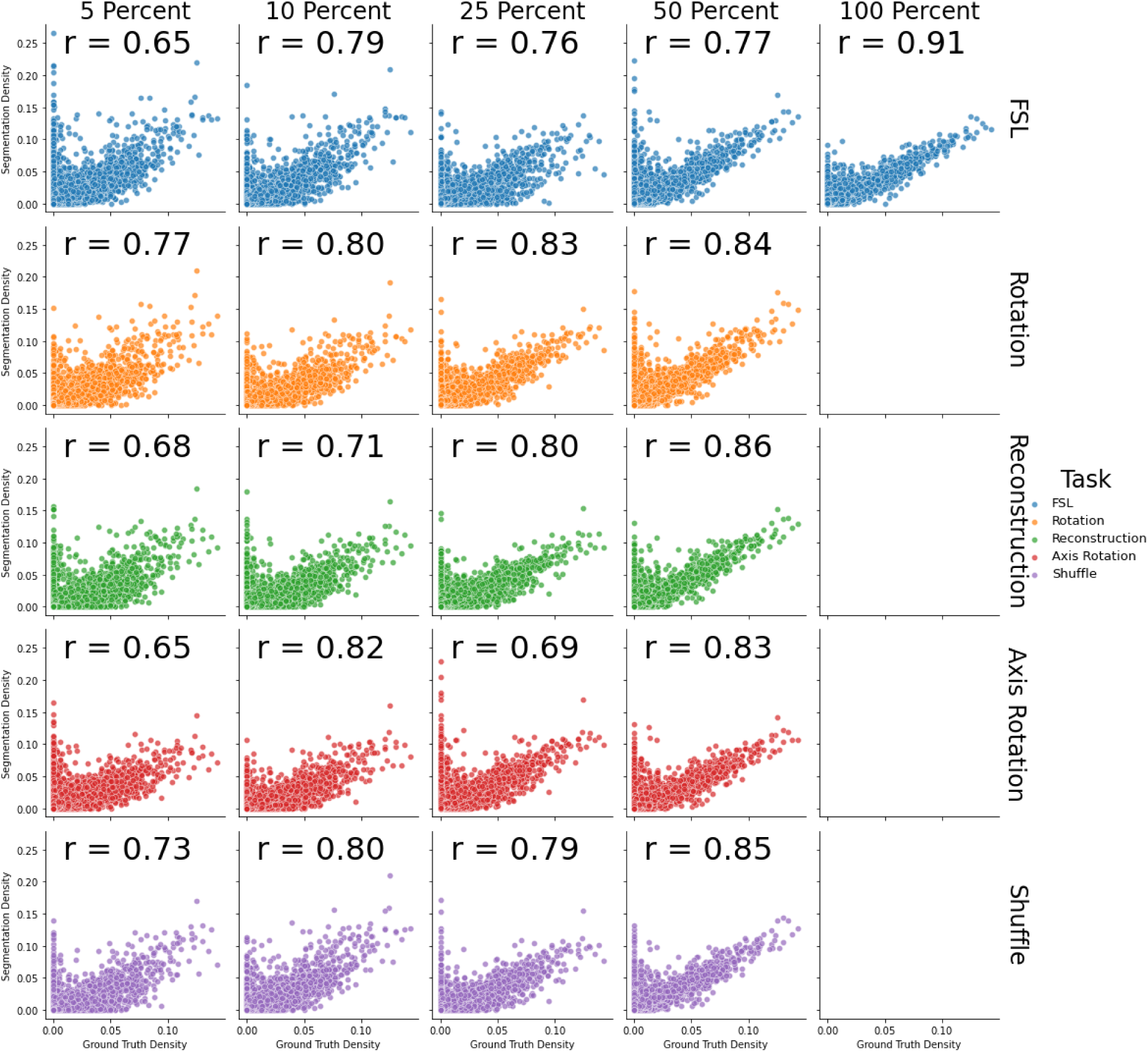
Scatterplot of segmentation neighborhood densities plotted against ground truth density from the neuron dataset. Volumes were divided into regions of size (32, 32, 32). Each row is a different method of training the model, each column is a percentage of training data used during finetuning. Inline: the Pearson R correlation coefficients between the pooled segmentations and the pooled ground truth

**Suppl Figure 5.**
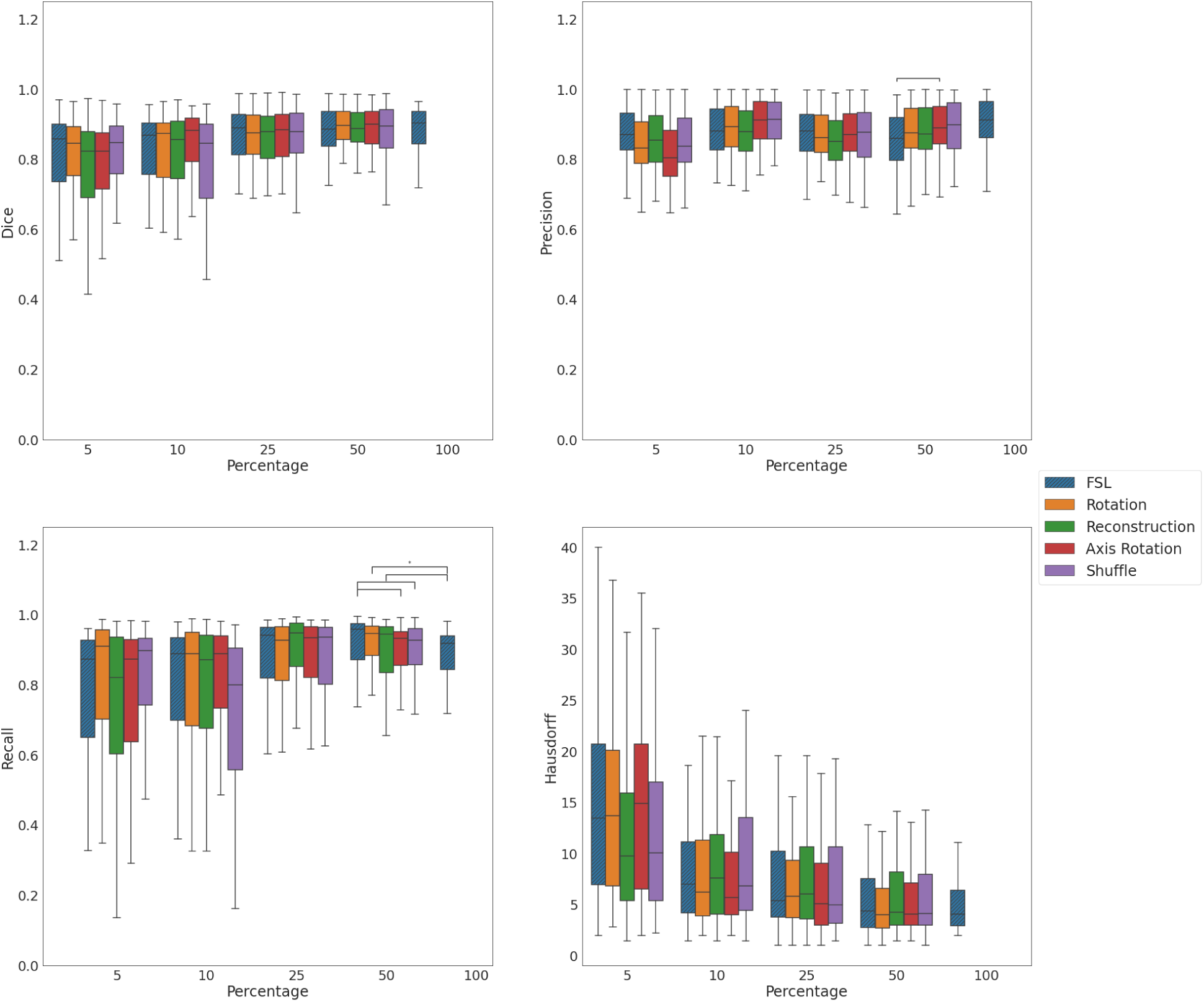
Boxplots of Dice coefficient, Precision, Recall, and Hausdorff Distances between ground truth vessel labels and segmentation masks generated by the FSL models and SSL models pretrained on all pretext tasks.

**Suppl Figure 6.**
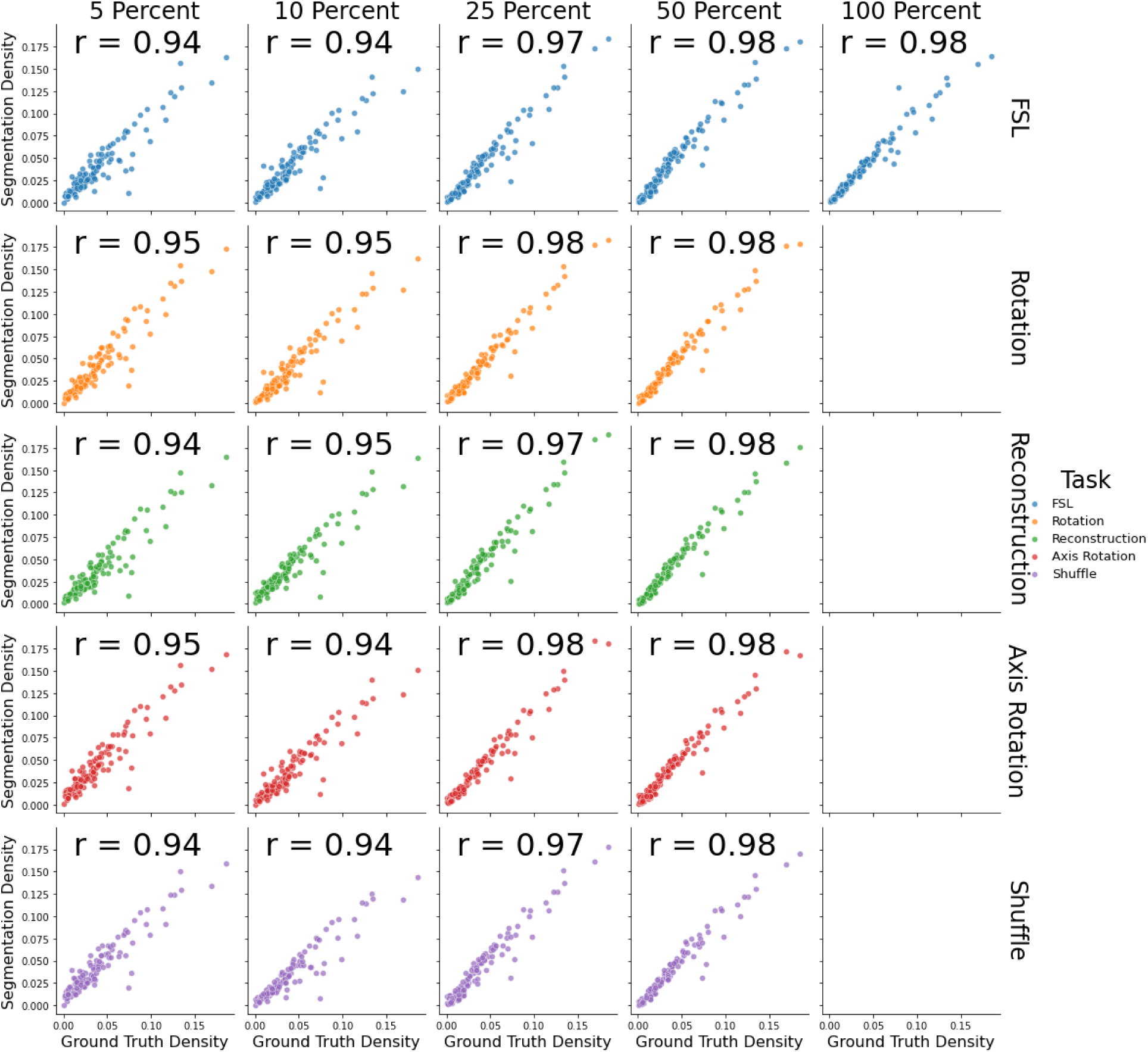
Scatterplot of segmentation neighborhood densities plotted against ground truth density. Each row is a different method of training the model, each column is a percentage of training data used during finetuning. Inline: the Pearson R correlation coefficients between the pooled segmentations and the pooled ground truth

**Suppl Figure 7.**
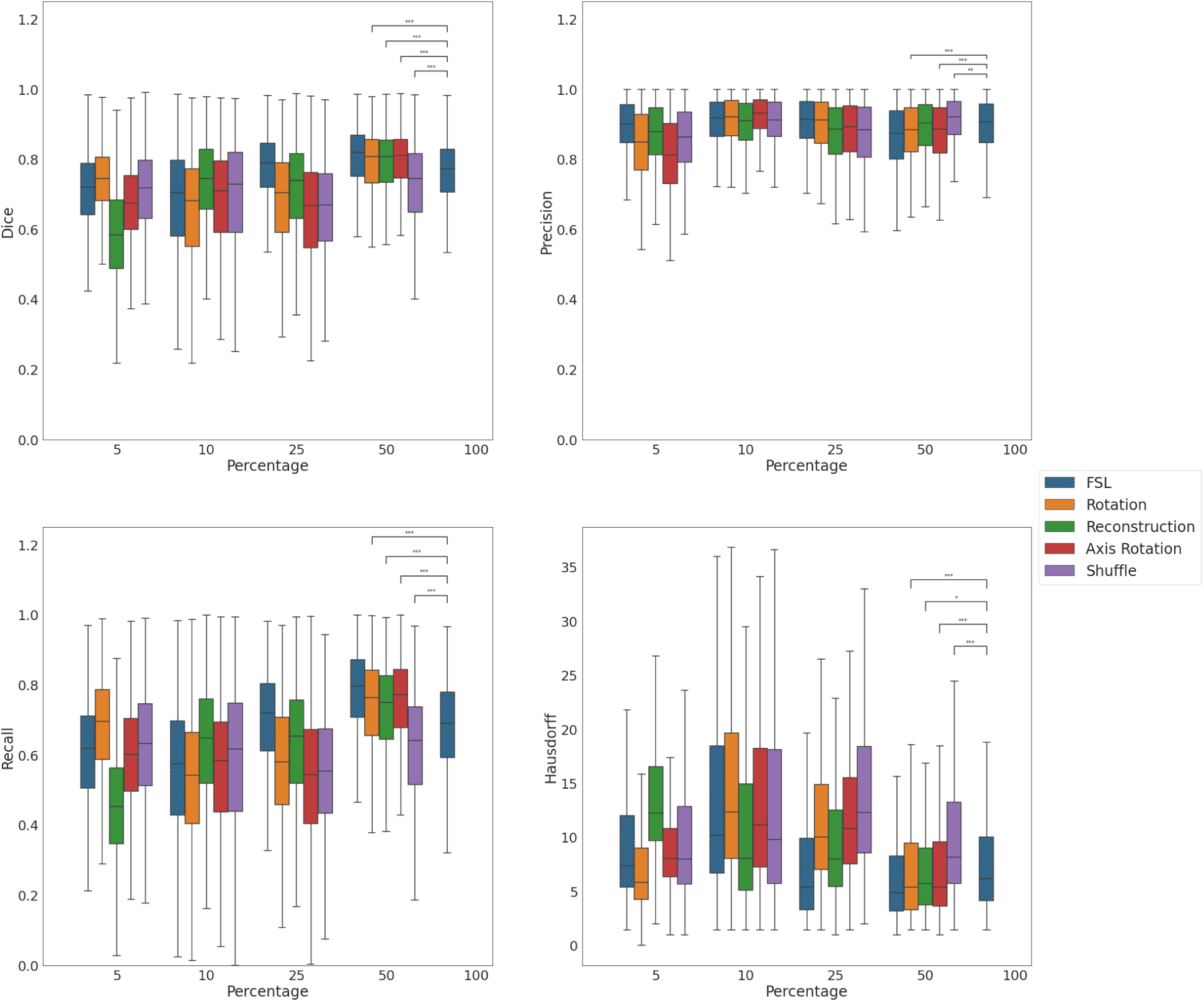
Boxplots of Dice coefficient, Precision, Recall, and Hausdorff Distances between MiniVess dataset ground truth and segmentation masks generated by the FSL models and SSL models pretrained on all pretext tasks.

**Suppl Figure 8.**
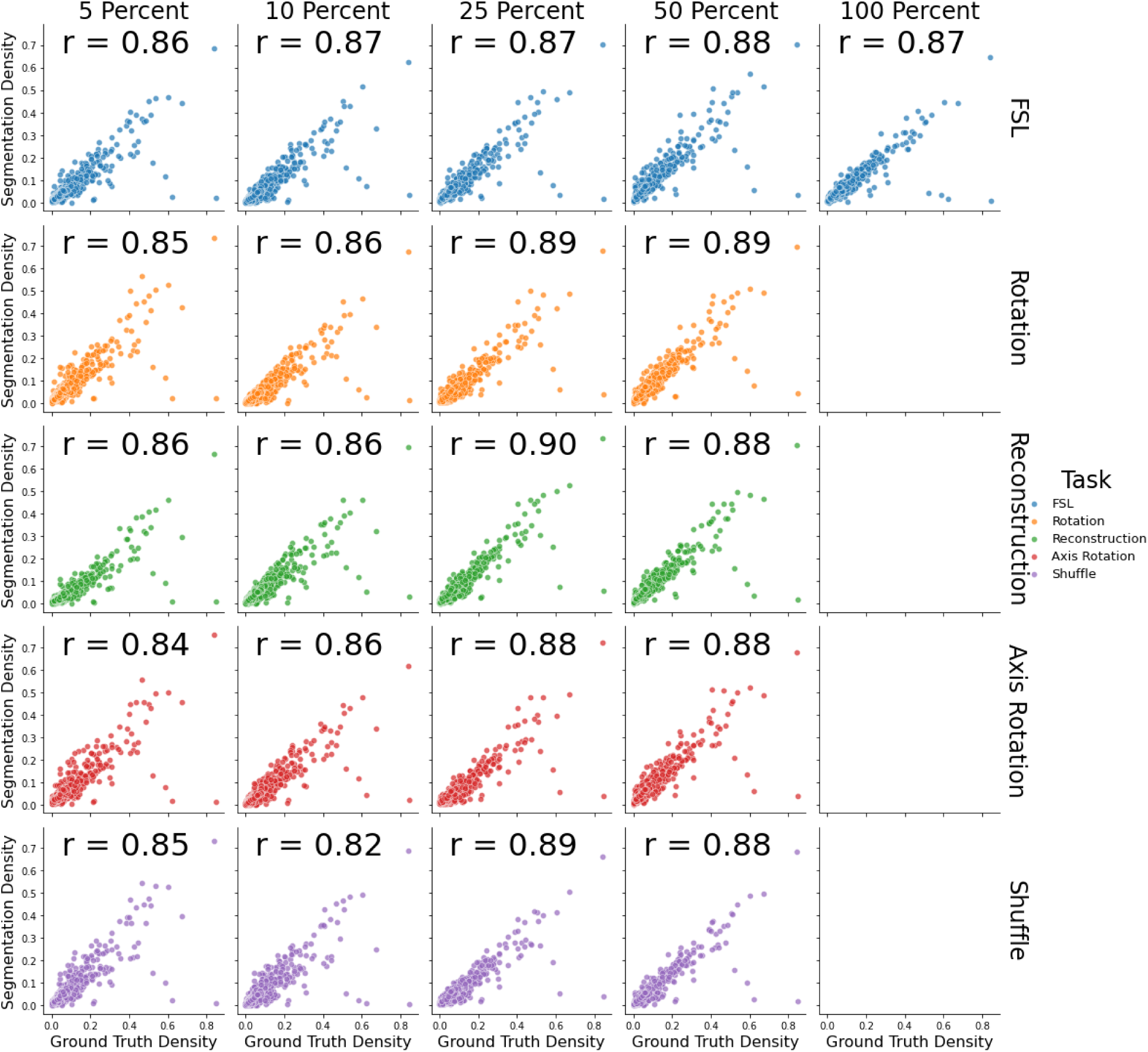
Scatterplot of segmentation neighborhood densities plotted against ground truth density. Each row is a different method of training the model, each column is a percentage of training data used during finetuning. Inline: the Pearson R correlation coefficients between the pooled segmentations and the pooled ground truth

**Suppl Figure 9.**
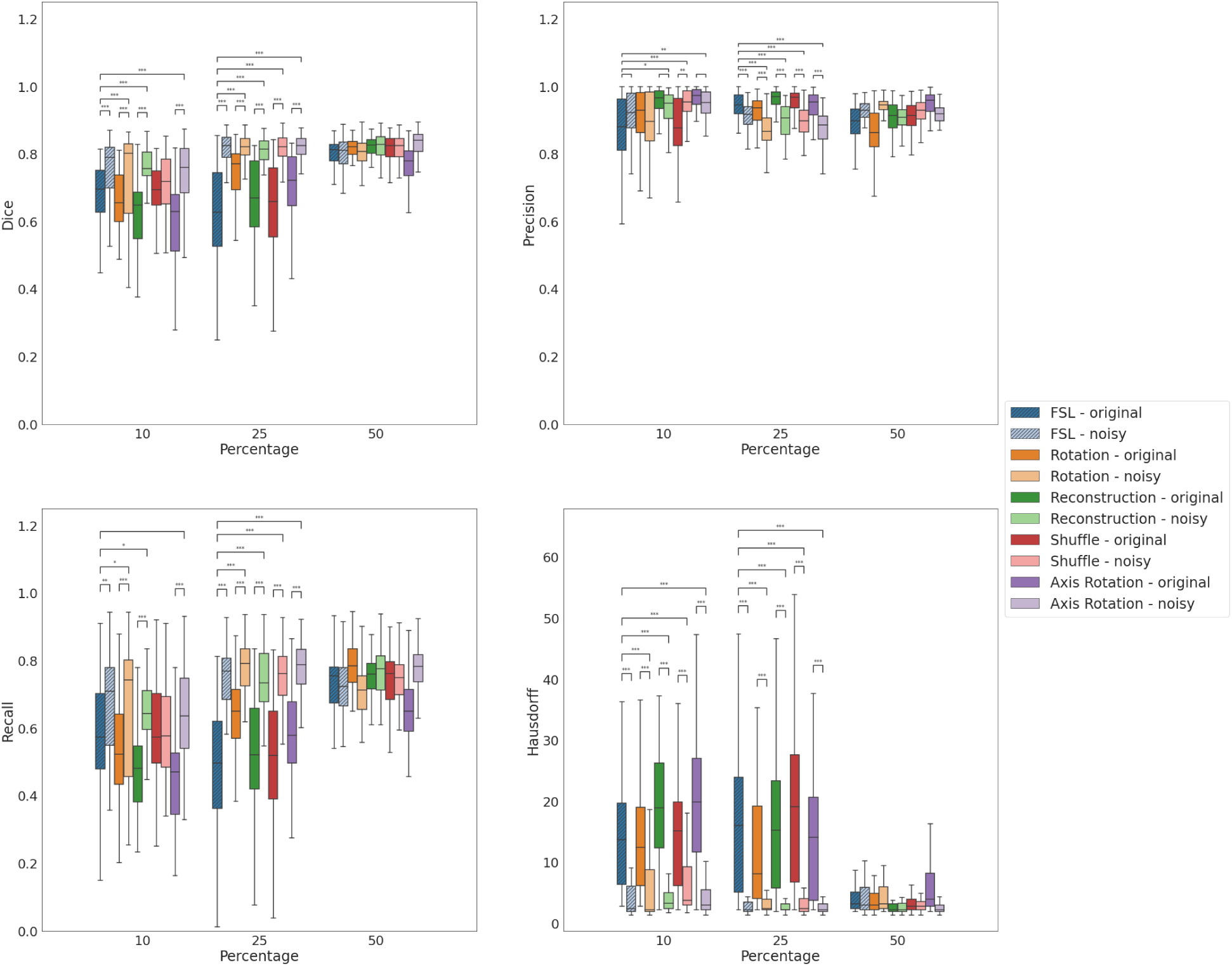
Boxplots of Dice coefficient, Precision, Recall, and Hausdorff Distances between ground truth neuron labels and segmentation masks generated by the FSL models and SSL models pretrained on all pretext tasks. Lighter shades designated for models that were finetuned with noisy datasets.

**Suppl Figure 10.**
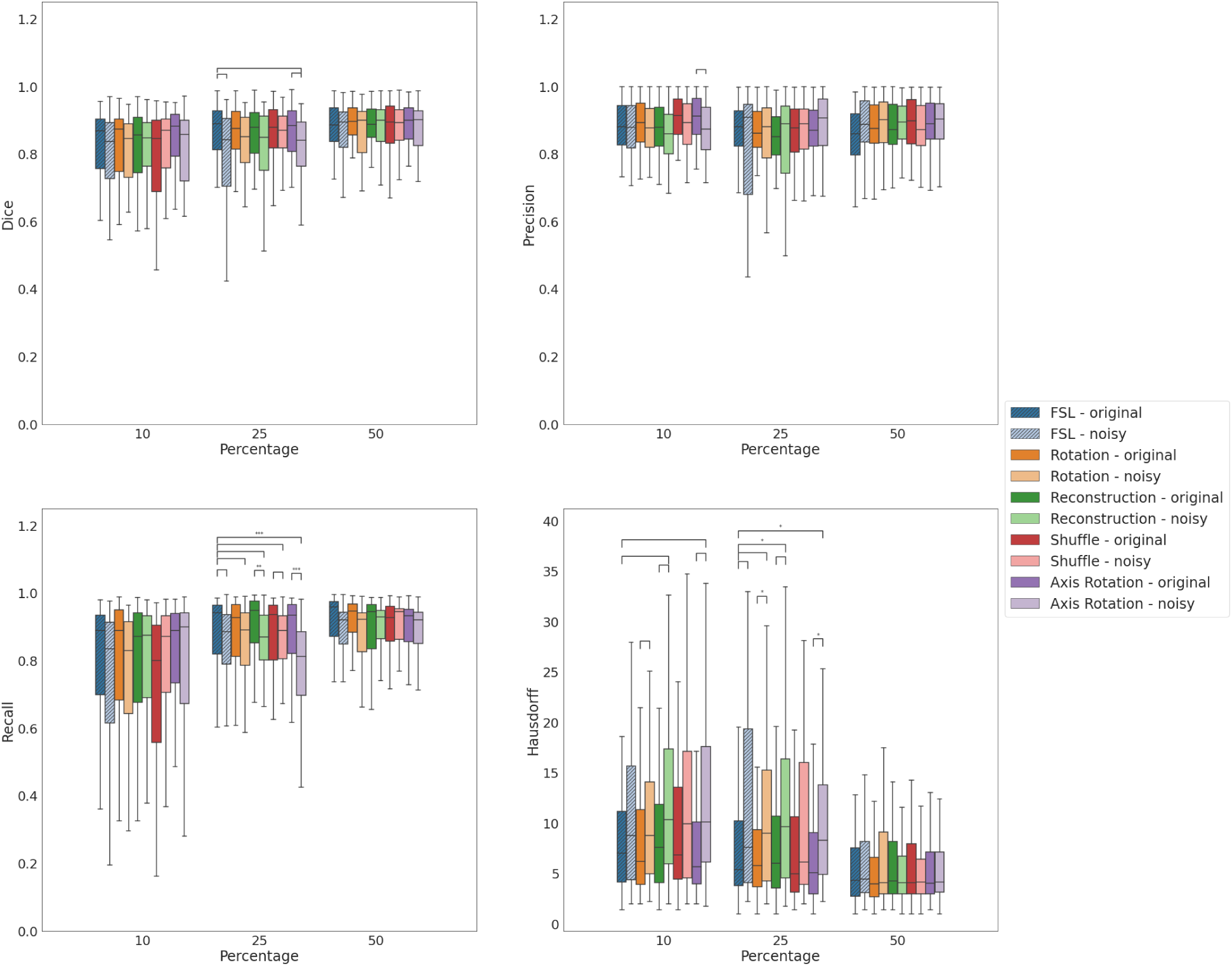
Boxplots of Dice coefficient, Precision, Recall, and Hausdorff Distances between ground truth vessel labels and segmentation masks generated by the FSL models and SSL models pretrained on all pretext tasks. Lighter shades designated for models that were finetuned with noisy datasets.

**Suppl Figure 11.**
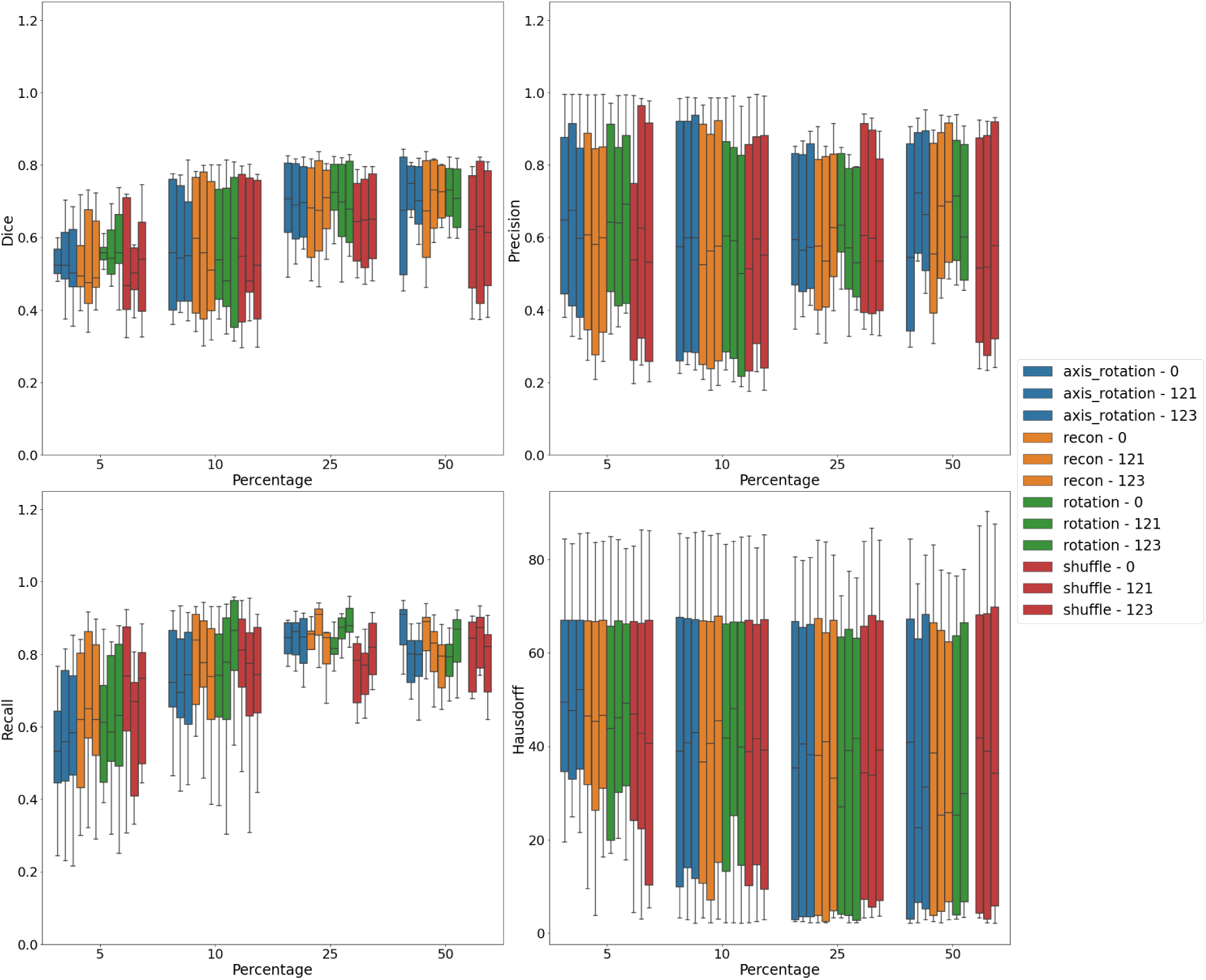
Boxplots of Dice coefficient, Precision, Recall, and Hausdorff Distances between ground truth neuron labels and segmentation masks generated by and SSL models finetuned with different seed initializations

**Suppl Figure 12.**
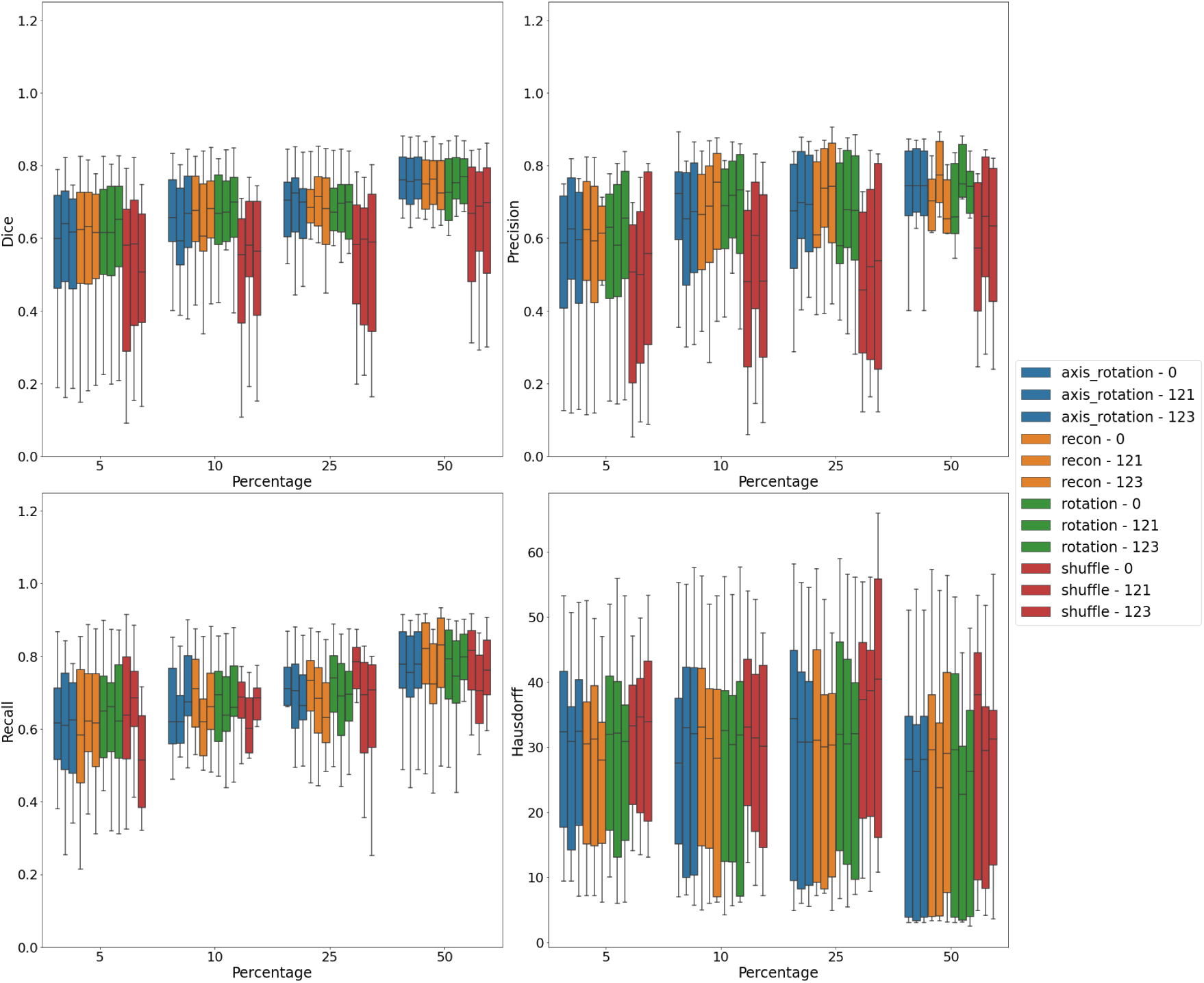
Boxplots of Dice coefficient, Precision, Recall, and Hausdorff Distances between ground truth vessel labels and segmentation masks generated by and SSL models finetuned with different seed initializations.

**Suppl Fig 13.**
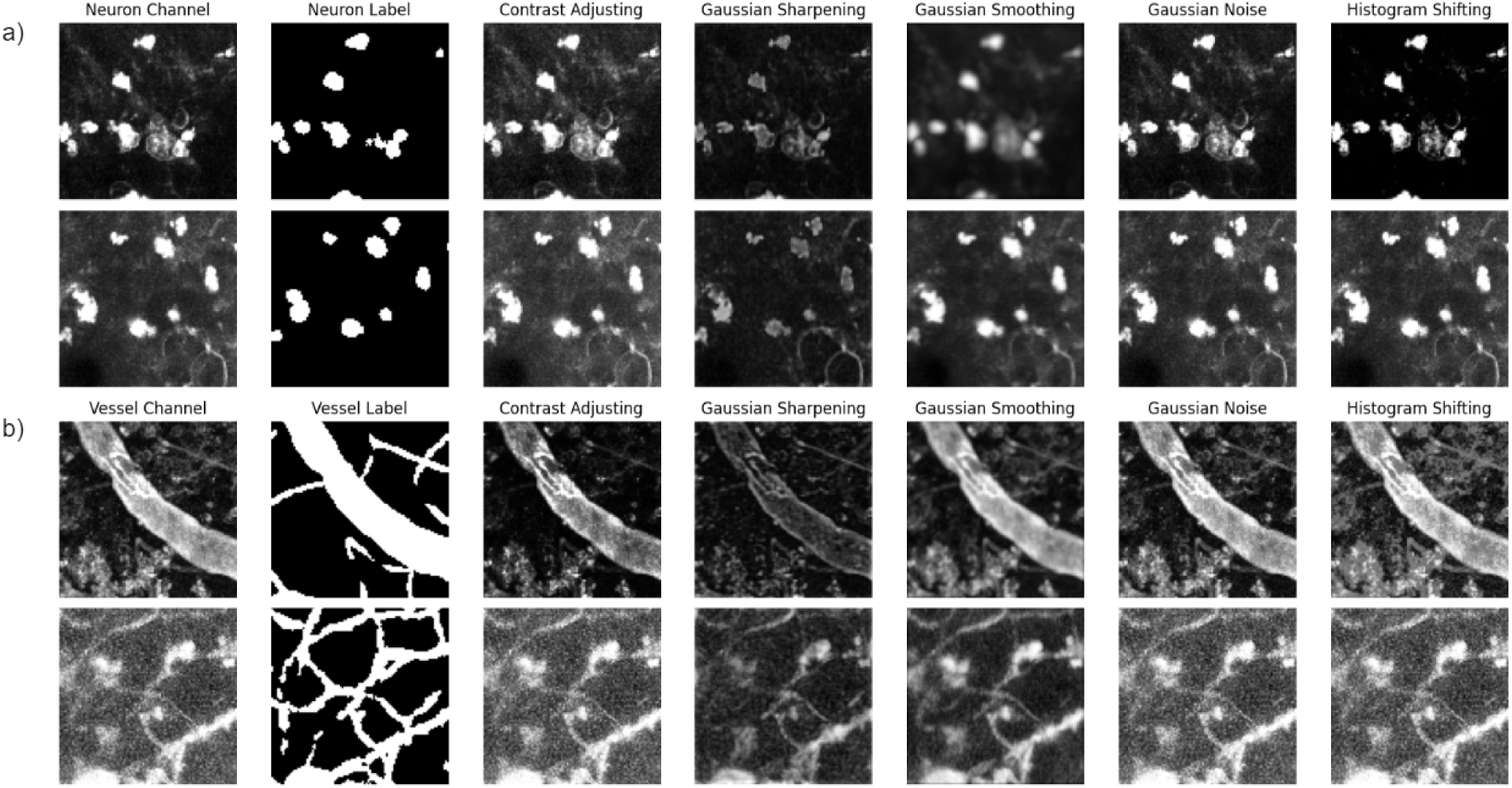
Example patch, and transforms used to finetune SSL and FSL models. In this figure, transforms are applied to either the neuron channel (a) or vessel channel (b).

**Suppl Table 1:** List of hyperparameters used in experiments for SSL pretraining.

|  | Hyperparameters | Shuffle | Rotation | Axis Rotation | Reconstruction |
| --- | --- | --- | --- | --- | --- |
| Pretraining | Epochs | 1000; early stopping at 100 epochs | 1000; early stopping at 100 epochs | 1000; early stopping at 100 epochs | 1000; early stopping at 100 epochs |
|  | Batch size | 64 | 64 | 64 | 64 |
|  | Learning Rate | Neuron: 0.001, Vessel: 1e-4 | Neuron: 0.01, Vessel: 1e-4 | Neuron: 0.001, Vessel: 1e-4 | Neuron: 0.0001, Vessel: 1e-4 |
|  | Optimizer | Adam | Adam | Adam | Adam |
|  | Scheduler | Multistep with lr decay at [300, 600] epochs by 0.1 | Multistep with lr decay at [300, 600] epochs by 0.1 | Multistep with lr decay at [300, 600] epochs by 0.1 | Multistep with lr decay at [300, 600] epochs by 0.1 |
|  | Selection of best model | Highest validation accuracy, lowest validation loss | Highest validation accuracy, lowest validation loss | Highest validation accuracy, lowest validation loss | Lowest validation loss |

**Suppl Table 2:** Mean scores for different models when segmenting the test neuron dataset.

| Percentage | Task | Dice | Precision | Recall | Hausdorff |
| --- | --- | --- | --- | --- | --- |
| 5 | FSL | **0.687 ± 0.081** | 0.849 ± 0.125 | 0.615 ± 0.157 | **11.532 ± 9.075** |
|  | Axis Rotation | 0.621 ± 0.12 | **0.958 ± 0.044** | 0.475 ± 0.135 | 13.541 ± 9.11 |
|  | Reconstruction | 0.567 ± 0.138 | 0.898 ± 0.095 | 0.446 ± 0.173 | 25.71 ± 12.606 |
|  | Rotation | 0.677 ± 0.112 | 0.861 ± 0.114 | **0.601 ± 0.186** | 12.635 ± 9.866 |
|  | Shuffle | 0.538 ± 0.165 | 0.94 ± 0.067 | 0.406 ± 0.185 | 27.686 ± 13.887 |
| 10 | FSL | 0.677 ± 0.101 | 0.871 ± 0.104 | 0.586 ± 0.161 | 14.933 ± 9.143 |
|  | Axis Rotation | 0.597 ± 0.138 | **0.957 ± 0.048** | 0.453 ± 0.152 | 20.819 ± 11.763 |
|  | Reconstruction | 0.62 ± 0.126 | 0.948 ± 0.055 | 0.481 ± 0.154 | 19.675 ± 11.701 |
|  | Rotation | 0.661 ± 0.094 | 0.911 ± 0.081 | 0.54 ± 0.139 | **13.638 ± 9.001** |
|  | Shuffle | **0.691 ± 0.083** | 0.876 ± 0.1 | **0.598 ± 0.149** | 14.771 ± 9.425 |
| 25 | FSL | 0.601 ± 0.201 | 0.936 ± 0.051 | 0.477 ± 0.205 | 16.66 ± 11.596 |
|  | Axis Rotation | 0.691 ± 0.122 | 0.935 ± 0.053 | 0.566 ± 0.144 | 13.186 ± 9.588 |
|  | Reconstruction | 0.652 ± 0.141 | **0.955 ± 0.045** | 0.516 ± 0.161 | 15.976 ± 11.323 |
|  | Rotation | **0.733 ± 0.099** | 0.923 ± 0.053 | **0.624 ± 0.132** | **11.696 ± 8.741** |
|  | Shuffle | 0.634 ± 0.162 | 0.954 ± 0.041 | 0.498 ± 0.172 | 19.818 ± 13.843 |
| 50 | FSL | 0.799 ± 0.044 | 0.886 ± 0.061 | 0.736 ± 0.08 | 5.571 ± 5.472 |
|  | Axis Rotation | 0.769 ± 0.058 | **0.945 ± 0.046** | 0.658 ± 0.096 | 7.693 ± 7.562 |
|  | Reconstruction | **0.816 ± 0.044** | 0.905 ± 0.052 | 0.748 ± 0.075 | **4.425 ± 5.118** |
|  | Rotation | 0.813 ± 0.043 | 0.864 ± 0.075 | **0.78 ± 0.086** | 5.215 ± 6.149 |
|  | Shuffle | 0.812 ± 0.05 | 0.909 ± 0.049 | 0.742 ± 0.087 | 4.532 ± 5.192 |
| 100 | FSL | 0.794 ± 0.057 | 0.926 ± 0.045 | 0.702 ± 0.086 | 6.143 ± 6.065 |

**Suppl Table 3:**
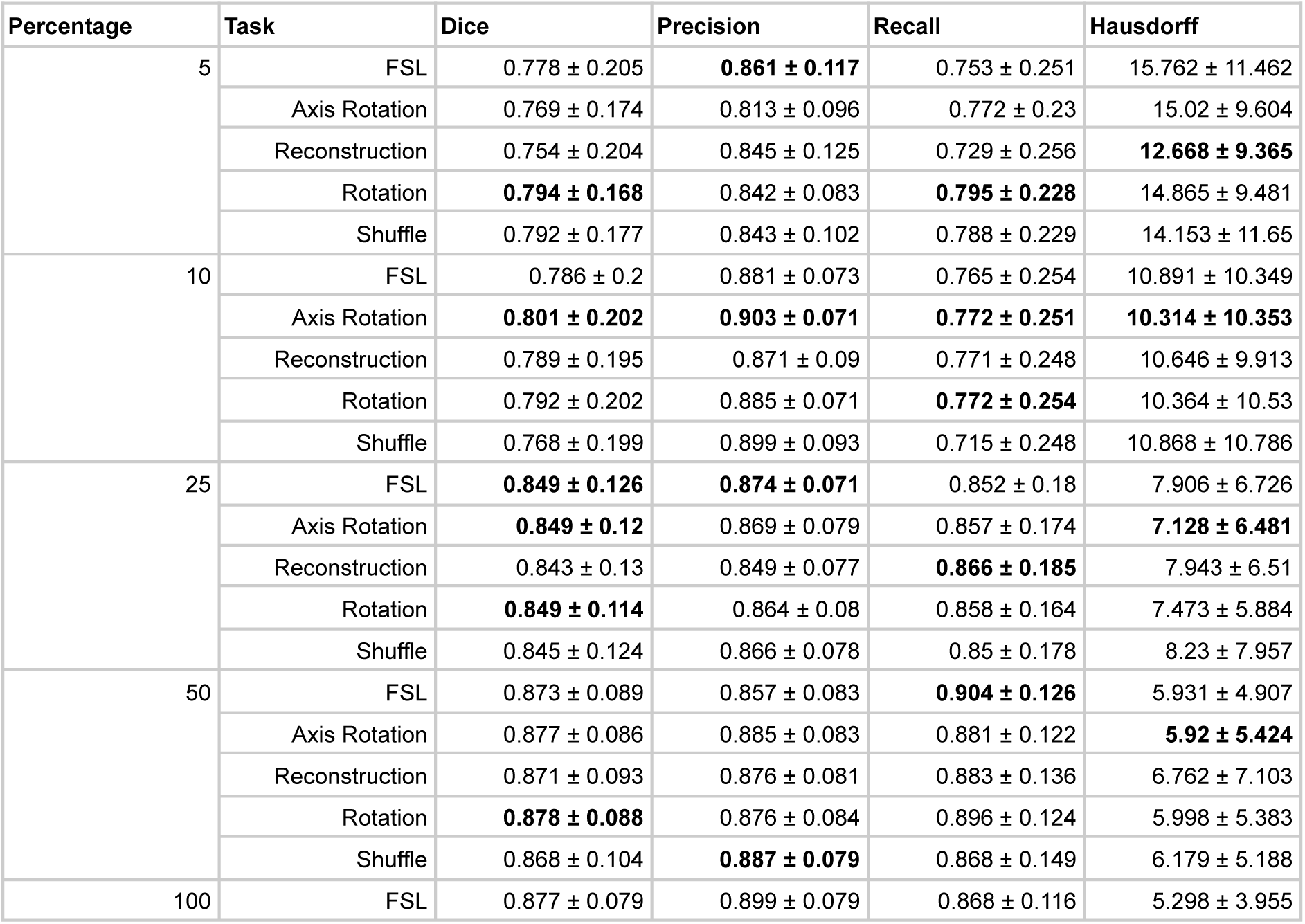
Mean scores for different models when segmenting the test vessels dataset.

**Suppl Table 4:** Mean scores for different models when segmenting the MiniVess dataset.

| Percentage | Task | Dice | Precision | Recall | Hausdorff |
| --- | --- | --- | --- | --- | --- |
| 5 | FSL | 0.698 ± 0.15 | **0.892 ± 0.08** | 0.598 ± 0.172 | 10.45 ± 9.123 |
|  | Axis Rotation | 0.659 ± 0.152 | 0.811 ± 0.116 | 0.585 ± 0.176 | 10.67 ± 9.329 |
|  | Reconstruction | 0.578 ± 0.158 | 0.87 ± 0.097 | 0.453 ± 0.163 | 14.657 ± 9.289 |
|  | Rotation | **0.727 ± 0.141** | 0.842 ± 0.103 | **0.671 ± 0.174** | **8.68 ± 8.664** |
|  | Shuffle | 0.694 ± 0.164 | 0.855 ± 0.101 | 0.616 ± 0.196 | 10.71 ± 8.871 |
| 10 | FSL | 0.661 ± 0.199 | 0.9 ± 0.1 | 0.554 ± 0.215 | 14.08 ± 10.984 |
|  | Axis Rotation | 0.677 ± 0.179 | **0.919 ± 0.077** | 0.564 ± 0.198 | 14.561 ± 11.18 |
|  | Reconstruction | **0.715 ± 0.168** | 0.898 ± 0.079 | **0.623 ± 0.194** | **11.699 ± 10.447** |
|  | Rotation | 0.64 ± 0.195 | 0.903 ± 0.099 | 0.525 ± 0.206 | 15.73 ± 11.496 |
|  | Shuffle | 0.679 ± 0.196 | 0.901 ± 0.088 | 0.581 ± 0.22 | 13.499 ± 11.376 |
| 25 | FSL | **0.765 ± 0.135** | **0.898 ± 0.084** | **0.691 ± 0.167** | **8.224 ± 8.236** |
|  | Axis Rotation | 0.643 ± 0.169 | 0.875 ± 0.1 | 0.536 ± 0.192 | 12.725 ± 8.226 |
|  | Reconstruction | 0.712 ± 0.152 | 0.868 ± 0.101 | 0.63 ± 0.183 | 10.381 ± 7.858 |
|  | Rotation | 0.681 ± 0.153 | 0.891 ± 0.092 | 0.576 ± 0.181 | 12.066 ± 8.367 |
|  | Shuffle | 0.646 ± 0.166 | 0.865 ± 0.107 | 0.542 ± 0.184 | 14.97 ± 9.721 |
| 50 | FSL | 0.768 ± 0.139 | 0.803 ± 0.116 | **0.781 ± 0.185** | **8.114 ± 9.682** |
|  | Axis Rotation | **0.785 ± 0.124** | 0.871 ± 0.098 | 0.74 ± 0.16 | 8.215 ± 8.314 |
|  | Reconstruction | 0.783 ± 0.119 | 0.887 ± 0.091 | 0.723 ± 0.153 | 8.144 ± 8.38 |
|  | Rotation | 0.778 ± 0.132 | 0.869 ± 0.099 | 0.73 ± 0.166 | 8.142 ± 8.531 |
|  | Shuffle | 0.711 ± 0.158 | **0.903 ± 0.082** | 0.612 ± 0.182 | 11.25 ± 9.375 |
| 100 | FSL | 0.743 ± 0.143 | 0.889 ± 0.092 | 0.665 ± 0.168 | 9.013 ± 9.482 |

**Suppl Table 5:** Mean scores for different models when segmenting the neuron dataset after finetuning/training with original and noisy datasets.

| Percentage | Task | Finetuning Data | Dice | Precision | Recall | HD95 |
| --- | --- | --- | --- | --- | --- | --- |
| 10 | FSL | original | 0.677 ± 0.101 | 0.871 ± 0.104 | 0.586 ± 0.161 | 14.933 ± 9.143 |
|  |  | noisy | 0.757 ± 0.086 | 0.909 ± 0.087 | **0.675 ± 0.149** | 7.122 ± 9.096 |
|  | Reconstruction | original | 0.62 ± 0.126 | **0.948 ± 0.055** | 0.481 ± 0.154 | 19.675 ± 11.701 |
|  |  | noisy | **0.761 ± 0.054** | 0.927 ± 0.075 | 0.66 ± 0.11 | **6.396 ± 7.418** |
|  | Rotation | original | 0.661 ± 0.094 | 0.911 ± 0.081 | 0.54 ± 0.139 | 13.638 ± 9.001 |
|  |  | noisy | 0.738 ± 0.129 | 0.892 ± 0.094 | 0.673 ± 0.197 | 8.312 ± 11.364 |
| 25 | FSL | original | 0.601 ± 0.201 | 0.936 ± 0.051 | 0.477 ± 0.205 | 16.66 ± 11.596 |
|  |  | noisy | **0.817 ± 0.042** | 0.9 ± 0.06 | 0.758 ± 0.087 | 4.201 ± 5.103 |
|  | Reconstruction | original | 0.652 ± 0.141 | **0.955 ± 0.045** | 0.516 ± 0.161 | 15.976 ± 11.323 |
|  |  | noisy | 0.81 ± 0.039 | 0.893 ± 0.065 | 0.753 ± 0.094 | **3.912 ± 4.958** |
|  | Rotation | original | 0.733 ± 0.099 | 0.923 ± 0.053 | 0.624 ± 0.132 | 11.696 ± 8.741 |
|  |  | noisy | 0.816 ± 0.043 | 0.865 ± 0.06 | **0.782 ± 0.088** | 4.919 ± 5.94 |
| 50 | FSL | original | 0.799 ± 0.044 | 0.886 ± 0.061 | 0.736 ± 0.08 | 5.571 ± 5.472 |
|  |  | noisy | 0.799 ± 0.057 | 0.919 ± 0.051 | 0.717 ± 0.098 | 5.666 ± 6.031 |
|  | Reconstruction | original | 0.816 ± 0.044 | 0.905 ± 0.052 | 0.748 ± 0.075 | **4.425 ± 5.118** |
|  |  | noisy | **0.823 ± 0.044** | 0.896 ± 0.056 | 0.768 ± 0.081 | 4.643 ± 6.013 |
|  | Rotation | original | 0.813 ± 0.043 | 0.864 ± 0.075 | **0.78 ± 0.086** | 5.215 ± 6.149 |
|  |  | noisy | 0.798 ± 0.055 | **0.934 ± 0.047** | 0.705 ± 0.093 | 6.012 ± 6.552 |

**Suppl Table 6:** Mean scores for different models when segmenting the vessel dataset after finetuning/training with original and noisy datasets.

| Percentage | Task | Finetuning Data | Dice | Precision | Recall | Hausdorff |
| --- | --- | --- | --- | --- | --- | --- |
| 10 | FSL | original | $0.786 \pm 0.2$ | **$0.881 \pm 0.073$** | $0.765 \pm 0.254$ | $10.891 \pm 10.349$ |
| | | noisy | $0.767 \pm 0.197$ | $0.871 \pm 0.102$ | $0.732 \pm 0.247$ | $12.119 \pm 9.585$ |
| | Reconstruction | original | $0.789 \pm 0.195$ | $0.871 \pm 0.09$ | $0.771 \pm 0.248$ | $10.646 \pm 9.913$ |
| | | noisy | **$0.794 \pm 0.166$** | $0.861 \pm 0.087$ | **$0.777 \pm 0.223$** | $13.557 \pm 9.517$ |
| | Rotation | original | $0.792 \pm 0.202$ | $0.885 \pm 0.071$ | $0.772 \pm 0.254$ | **$10.364 \pm 10.53$** |
| | | noisy | $0.769 \pm 0.194$ | $0.876 \pm 0.088$ | $0.732 \pm 0.244$ | $12.238 \pm 9.998$ |
| 25 | FSL | original | **$0.849 \pm 0.126$** | **$0.874 \pm 0.071$** | $0.852 \pm 0.18$ | $7.906 \pm 6.726$ |
| | | noisy | $0.787 \pm 0.162$ | $0.826 \pm 0.171$ | $0.812 \pm 0.199$ | $12.048 \pm 9.004$ |
| | Reconstruction | original | $0.843 \pm 0.13$ | $0.849 \pm 0.077$ | **$0.866 \pm 0.185$** | $7.943 \pm 6.51$ |
| | | noisy | $0.81 \pm 0.139$ | $0.843 \pm 0.146$ | $0.818 \pm 0.179$ | $11.887 \pm 9.298$ |
| | Rotation | original | $0.849 \pm 0.114$ | $0.864 \pm 0.08$ | $0.858 \pm 0.164$ | **$7.473 \pm 5.884$** |
| | | noisy | $0.822 \pm 0.121$ | $0.834 \pm 0.15$ | $0.84 \pm 0.149$ | $11.444 \pm 8.886$ |
| 50 | FSL | original | $0.873 \pm 0.089$ | $0.857 \pm 0.083$ | **$0.904 \pm 0.126$** | **$5.931 \pm 4.907$** |
| | | noisy | $0.857 \pm 0.112$ | **$0.88 \pm 0.088$** | $0.858 \pm 0.157$ | $6.883 \pm 6.014$ |
| | Reconstruction | original | $0.871 \pm 0.093$ | $0.876 \pm 0.081$ | $0.883 \pm 0.136$ | $6.762 \pm 7.103$ |
| | | noisy | $0.872 \pm 0.089$ | $0.88 \pm 0.084$ | $0.879 \pm 0.125$ | $5.709 \pm 5.179$ |
| | Rotation | original | **$0.878 \pm 0.088$** | $0.876 \pm 0.084$ | $0.896 \pm 0.124$ | $5.998 \pm 5.383$ |
| | | noisy | $0.853 \pm 0.113$ | $0.878 \pm 0.096$ | $0.849 \pm 0.156$ | $6.727 \pm 5.552$ |

**Suppl Table 7:** Mean scores for different models when segmenting the MiniVess dataset after finetuning/training with original and noisy datasets.

| Percentage | Task | Finetuning Data | Dice | Precision | Recall | Hausdorff |
| --- | --- | --- | --- | --- | --- | --- |
| 10 | FSL | original | $0.659 \pm 0.2$ | $0.899 \pm 0.101$ | $0.552 \pm 0.217$ | $14.247 \pm 11.024$ |
| | | noisy | $0.581 \pm 0.166$ | $0.899 \pm 0.078$ | $0.451 \pm 0.167$ | $13.815 \pm 8.95$ |
| | Reconstruction | original | **$0.712 \pm 0.168$** | $0.897 \pm 0.08$ | **$0.62 \pm 0.195$** | $11.866 \pm 10.481$ |
| | | noisy | $0.67 \pm 0.145$ | $0.872 \pm 0.091$ | $0.567 \pm 0.165$ | **$10.734 \pm 8.402$** |
| | Rotation | original | $0.638 \pm 0.197$ | **$0.901 \pm 0.1$** | $0.523 \pm 0.208$ | $15.895 \pm 11.539$ |
| | | noisy | $0.692 \pm 0.149$ | $0.867 \pm 0.094$ | $0.602 \pm 0.171$ | $10.936 \pm 9.319$ |
| 25 | FSL | original | **$0.764 \pm 0.136$** | **$0.897 \pm 0.084$** | $0.69 \pm 0.169$ | **$8.325 \pm 8.285$** |
| | | noisy | $0.505 \pm 0.218$ | $0.866 \pm 0.129$ | $0.402 \pm 0.234$ | $21.304 \pm 9.772$ |
| | Reconstruction | original | $0.711 \pm 0.153$ | $0.866 \pm 0.101$ | $0.628 \pm 0.185$ | $10.51 \pm 7.881$ |
| | | noisy | $0.681 \pm 0.139$ | $0.789 \pm 0.142$ | **$0.641 \pm 0.189$** | $13.584 \pm 9.474$ |
| | Rotation | original | $0.68 \pm 0.154$ | $0.89 \pm 0.092$ | $0.575 \pm 0.182$ | $12.184 \pm 8.406$ |
| | | noisy | $0.663 \pm 0.148$ | $0.811 \pm 0.143$ | $0.596 \pm 0.191$ | $15.791 \pm 9.466$ |
| 50 | FSL | original | **0.796 ± 0.116** | 0.858 ± 0.099 | **0.768 ± 0.154** | **7.521 ± 8.033** |
|  |  | noisy | 0.732 ± 0.123 | **0.915 ± 0.077** | 0.627 ± 0.15 | 12.03 ± 9.138 |
|  | Reconstruction | original | 0.782 ± 0.12 | 0.886 ± 0.091 | 0.722 ± 0.154 | 8.241 ± 8.433 |
|  |  | noisy | 0.763 ± 0.125 | 0.866 ± 0.102 | 0.707 ± 0.161 | 8.442 ± 7.95 |
|  | Rotation | original | 0.776 ± 0.133 | 0.868 ± 0.099 | 0.729 ± 0.167 | 8.24 ± 8.586 |
|  |  | noisy | 0.761 ± 0.131 | 0.888 ± 0.094 | 0.691 ± 0.164 | 9.456 ± 9.724 |

## Notes

https://search.kg.ebrains.eu/instances/bf268b89-1420-476b-b428-b85a913eb523

### Competing Interest Statement

The authors have declared no competing interest.

